# LipiDetective a deep learning model for the identification of molecular lipid species in tandem mass spectra

**DOI:** 10.1101/2024.10.07.617094

**Authors:** Vivian Würf, Nikolai Köhler, Florian Molnar, Lisa Hahnefeld, Robert Gurke, Michael Witting, Josch K. Pauling

## Abstract

Lipids are involved in many vital processes within the cell, and alterations in lipid homeostasis have been associated with various diseases such as cancer or type 2 diabetes. Confidently identifying lipids in samples is a prerequisite for understanding the multiple functions lipids fulfill in health and disease. However, the accurate identification of molecular lipid species based on tandem mass spectrometry data is still a key challenge in lipidomics. Most current approaches rely on using a custom pipeline to process and match the measured spectra against an in-house spectra reference library, which hinders the comparability of results. To address this challenge, a transformer model called LipiDetective was developed and trained on a dataset composed of reference spectra measured from lipid standards, spectra from databases, and published experiments, utilizing both shotgun as well as liquid-chromatography mass spectrometry. LipiDetective demonstrates, for the first time, that artificial neural networks can learn the characteristic lipid fragmentation patterns to automatically and accurately annotate molecular lipids species in tandem mass spectra independently of the experimental setup. The model can even correctly predict lipid species for which it has never seen a spectrum before as it is able to generalize the learned lipid fragmentation patterns. Analysis of the integrated gradients reveals that LipiDetective focuses on relevant peaks that can be matched to known fragments and are thus humanly interpretable. Therefore, LipiDetective has the potential to be a valuable tool to aid in the lipid identification process and support the comparability of results from different sources. Aside from Lipidetective as a “ready-to-use” application, this work primarily offers a deeper understanding of how the model functions and how future deep learning models for lipid identification in mass spectra could be improved.

## 1 Introduction

Lipids have many important functions in biological systems; they serve as energy and heat sources, are involved in cell signaling and act as the main structural elements of biological membranes [1]. To fulfill all these roles, they exhibit an enormous structural variety, with mammalian cells producing thousands of different lipid species and utilizing hundreds of proteins for their synthesis, remodeling, transport and degradation [2]. As of September 2024, the Lipid Maps Structure Database (LMSD) [3] contained 26,534 curated unique lipid structures. Technical advancements in mass spectrometry (MS) and bioinformatics enable a higher resolution than ever to identify and quantify the many lipid species contained within a single sample.

In a typical lipidomics workflow, the lipids from a biological sample are extracted and then analyzed using MS. In the mass spectrometer, the extracted lipids are first ionized, then separated based on their mass-to-charge (m/z) ratio, and finally, their relative abundance is measured by the ion detector [4]. The resulting mass spectrum shows the m/z ratio on the x-axis and the abundance of the respective ion on the y-axis [4]. This process alone does not allow for the differentiation between isomeric lipid species (meaning having the same molecular formula, but different atom connectivity) and only enables the annotation of lipids on a sum species level. Lipid sum species level means that only the total number of carbons and double bonds of all fatty acyl chains attached to a lipid can be inferred. Additionally, since two structurally distinct lipids can have identical masses, a measured m/z peak may correspond to two different lipid species, possibly from different lipid classes. Even if two lipids have only similar, but not identical, masses, they may still be indistinguishable if the mass spectrometer lacks sufficient resolving power. To characterize lipidomes down to their molecular lipid species composition, meaning the acyl composition, tandem mass spectrometry (MS2) can be employed.

The technique of MS2 involves two mass analyzers. In data-dependent aquisition (DDA), a first MS1 measurement is conducted as previously explained, after which particularly prevalent ions of a specific m/z are selected based on set criteria for further tandem MS analysis. These so-called precursor ions are isolated and then fragmented by collision with a neutral gas such as nitrogen or argon in a process called collision-induced dissociation (CID) [4]. Another precursor selection method, called data-independet acquisition (DIA), involves selecting and fragmenting ions within a specified m/z range. However, due to the mixed spectra generated by DIA, this acquisition method is not relevant for this work. The degree of dissociation during the fragmentation step can be adjusted via the collision energy parameter, which indicates the kinetic energy the ions carry after an acceleration stage [5]. The ion fragments resulting from this dissociation are then recorded in the second step of MS2 [4]. Based on the fragments resulting from the dissociation, it is possible to infer structural information, thereby allowing to distinguish between isomeric ions with the same m/z value in the MS1 [5].

The identification of lipids in the spectra is commonly done using software that matches the detected fragment peaks to a database entry. Identifying and quantifying the large diversity of lipid species contained within one biological sample is still one of the main challenges in lipidomics research. Most approaches for identifying lipids in mass spectra currently rely on comparing the measured MS2 spectra to an in-house database of fragmentation spectra using a custom software pipeline [6]. Different software tools may use varying approaches for the preprocessing and identification steps, which, combined with the use of divergent spectral databases, leads to low reproducibility and comparability between labs. This issue is exacerbated by the many varying parameters in the experimental setup, such as collision energy, polarity mode, and type of mass spectrometer.

In 2017, the lipid composition of the human plasma standard reference material (SRM) 1950 was measured by 31 diverse laboratories, each employing their custom lipidomics workflow [7]. Across all laboratories a total of 1527 unique lipids were identified on a sum composition level. However, there were only 339 lipids that were reported by five or more laboratories. Considering that this comparison was conducted on a sum composition level one can safely assume that the difference between the results would be even more significant when scaling up to molecular lipid species level.

This means that new and innovative approaches must be explored to improve comparability between results. Here, deep learning (DL) offers a promising alternative to spectrum matching approaches. DL has already been applied to a variety of tasks in MS data processing, such as noise filtering, peak detection, and metabolite annotation [8]. Still, the application of DL in lipidomics is in its infancy due to various challenges such as low sample sizes, lack of interpretability, and general lack of sufficient reference, training, and validation data. Nevertheless, this shows the enormous potential of applying artificial neural networks (ANNs) to MS data and can serve as a blueprint for how to approach similar problems in lipidomics research.

The main argument for applying an ANN to the task of lipid identification from mass spectra is, that the neural network will be able to learn and abstract the characteristic fragmentation pattern of each lipid species. Ideally, the neural network is then able to identify lipids independent of collision energy or other experimental setups and even generalize the learned lipid fragmentation patterns to unseen lipid species. As annotations by a single software can often lead to a high rate of false positive identifications [9], a DL model that is able to identify molecular lipid species from fragment spectra in a fast, automated, and accurate manner can serve as an additional means of validation. Improving data quality and confidence in lipid identifications will enhance the comparability of results across different laboratories, thereby facilitating progress in the field of lipidomics.

This motivated the development of LipiDetective, a transformer model that can identify lipids in MS2 spectra on a molecular lipid species level. As previously mentioned, developing such DL models comes with unique challenges, many of which center around the data on which the model is trained. Sufficient high-quality data, in this case confidently identified spectra of molecular lipid species, has to be available. Unfortunately, this kind of data is still challenging to obtain, as it is not yet common practice in lipidomics to supply the raw data with a publication. Even if the raw data is available, it is often not possible to match the identifications back to the exact corresponding spectra. Spectrum IDs are rarely provided and the metadata between the raw measurement files and annotation tables might differ due to data preprocessing steps such as retention time alignment. The dataset for training LipiDetective consists of spectra generated from measurements of chemical reference standards at varying collision energies specifically for this project as well as raw data that was available in publications and databases. Even with this relatively small training dataset of 268,720 spectra, LipiDetective was able to learn lipid fragmentation patterns independently of the experimental setup. Model interpretability methods such as integrated gradients and visualization of attention weights show that LipiDetective learns to associate peaks with certain lipid sub-structures. This means that it does not have to rely on matching the overall spectra but can identify relevant lipid components from which it composes the final prediction. This allows it to even identify lipid species for which it has never previously encountered a spectrum. LipiDetective provides a first foundation for the community to build and expand on to further utilize the potential of DL for lipidomics.

## 2 Methods

### 2.1 Datasets

#### 2.1.1 GNPS Database

The database Global Natural Products Social Molecular Networking (GNPS) [10] was searched for publicly available datasets containing MS2 spectra of lipids. As of April 2024, there are three datasets available for download as JSON files that are focused on lipids:

The **PNNL** dataset is composed of 46,724 MS2 spectra from 1790 lipids measured by Thomas Metz’s group at the Pacific Northwest National Lab (PNNL) [11, 12]. The spectra were collected in positive and negative ionization mode using varying collision energy. The **HCE** dataset consists of 116 spectra measured in negative mode of lipids extracted from human corneal epithelium cells [13]. The **IOBA-NHC** dataset contains 197 MS2 spectra of lipids from human conjunctival cells (IOBA-NHC cell line) with 106 measured in negative and 91 in positive mode [14]. The HCE and IOBA-NHC libraries were generated by the Olivier Laprévote Lab from the Université de Paris.

#### 2.1.2 MITOMICS Dataset

The mitochondrial orphan protein multi-omic CRISPR screen (MITOMICS) dataset was generated to establish a functional compendium of human mitochondrial proteins [15]. For this purpose, the proteome, metabolome, and lipidome of more than 200 CRISPR-mediated HAP1 cell knockout lines were profiled using MS. The raw files and corresponding mzML files are available in the MassIVE [16] data repository under accession #MSV000086685. Across all cell lines, the publication identified 1,349 lipids belonging to over 30 lipid classes using the LipiDex software [17].

The identifications were matched to the corresponding spectra in the 882 lipidomics mzML files using the provided retention time and precursor masses. Additionally, LMSD was queried to retrieve the exact mass of the identified lipid. As the adduct was not provided with the identification, it had to be manually determined as it is a component of the prediction output. For this purpose, a list of possible adducts was compiled from the LipiDex instructions, and using the exact mass as well as the precursor mass, the corresponding adduct was determined.

#### 2.1.3 Thermo

The Thermo dataset consists of measurements from a study published in 2023 on the effects of different storage conditions on lipid stability in mice tissue homogenates [18]. Samples from multiple tissues, including the liver, spleen, kidney, and heart, were measured using a Thermo Orbitrap Exploris 480 and normalized, stepped high-energy collisional dissociation (HCD) with 15, 30, and 50 eV. The MS2 spectra acquisition was data-dependent and with a mass resolution of 15.000. The lipids were identified on a sum species level utilizing the Compound Discoverer 3.1 from Thermo Fisher Scientific with the LipidBlast VS68 positive and negative libraries.

Building on this sum species identification in the study, the Alex^123^ database [19] was used to match peaks in the MS2 spectra to known fragments to elucidate the fatty acid composition. At least three fragment peaks had to match to qualify as a possible identification. In the case where multiple isomeric lipid species were possible matches, the species whose matching peaks had the highest summed intensity was chosen.

#### 2.1.4 Phospholipid Standards

A reference dataset composed of single phospholipid standard measurements was generated for this study using shotgun MS2 with electrospray ionization (ESI) as an ionization technique. It contains MS2 spectra of 54 different phospholipid standards measured in positive and negative mode, except for phosphatidylglycerols (PGs), which were only measured in negative mode (Supplement Table S2). CID was applied using stepped collision energies. The same standards were measured using mass spectrometers from three different instruments: Agilent 6560 IM QTOF-MS, Sciex X500R QTOF-MS, and Bruker maXis UHR-TOF-MS. Lipid standards were obtained from Avanti Polar Lipids and dissolved in an appropriate solvent. Stock solutions were diluted in methanol/isopropanol/chloroform (1/1/1, v/v/v) with 7.5 mM ammonium formate. Lipid were infused using a syringe-pump at a flow of 500 μL/h. MS2 spectra were collected in an multiple reaction monitoring (MRM)-type acquisition with different collision energies. The m/z values of different adducts covered were isolated and fragmented with collision energies from 10 to 50 eV in 2.5 eV steps. Data was exported as .mzML using MSConvert [20].

### 2.2 Data Preparation

#### 2.2.1 Preprocessing

All profile spectra underwent smoothing, baseline correction, and peak picking, for which the pyOpenMS [21] library was used. Additionally, base peak normalization was performed for all spectra. This means that all peak intensities are divided by the intensity of the respective base peak, which is the peak with the highest intensity in the individual spectrum. Thus, the normalized intensities in every spectrum lie between zero and one. As the maximum precursor mass of the heaviest lipid in the dataset was around 1520 m/z, all MS2 spectra were trimmed to an m/z range of 50 - 1600 m/z. The varying nomenclatures in the different data sources were manually adjusted to match the updated lipid shorthand notation [22].

Isomers tend to coelute in most methods leading to mixed spectra. Additionally, the ionization efficacy changes based on chain length and number of double bonds. The combination of these two effects means that the sn-position of the fatty acid side chains cannot be determined reliably solely based on an MS2 spectrum generated via CID. Therefore, fatty acids were always listed in order of increasing carbon chain length. In case a lipid contained multiple fatty acids with the same carbon chain length, they were sorted based on their number of double bonds in ascending order.

#### 2.2.2 Noisy Spectra Filtering

An additional filtering step was performed for the phospholipid standard measurements to exclude particularly noisy spectra that either do not show any target lipid or have a very low signal-to-noise ratio (Supplement Figure S1). In general, the median intensity seems to remain similar between noisy and high-quality spectra, whereas the base peak tends to have a much lower intensity in the noisy spectra. This characteristic was used to estimate the noise level of a spectrum by dividing the median intensity by the base peak intensity. To generate the high-quality standards dataset, a threshold of 0.01 base peak normalized median intensity was chosen, and any spectra with a higher score were excluded.

#### 2.2.3 HDF5 Datasets

After preprocessing, all spectra were saved in a Hierarchical Data Format 5 (HDF5) file [23] using h5py [24]. This was done so that the preprocessing steps only had to be performed once to allow for easy and fast data reading during training. Each spectrum was saved as an HDF5 dataset with the corresponding molecular lipid species, adduct, and precursor mass as metadata attributes.

### 2.3 LipiDetective Architecture

LipiDetective is an adaptation of the transformer model [25] and approaches the problem of lipid identification as a sequence-to-sequence task. In this case, the input sequence is a list of the top n m/z values, ordered by decreasing intensity. The output sequence is the name of the molecular lipid species in shorthand notation, including the adduct present in the spectrum (Figure 2). LipiDetective was implemented by modifying the transformer module provided by PyTorch [26]. To prepare the spectrum as input for the model, the m/z value of each input peak is truncated to a user-specified decimal point and then matched to its corresponding embedding vector. The positional encoding is then generated. Since the peaks have been ordered by decreasing intensity, the positional encoding incorporates how high peaks are in relation to each other without relying on their explicit values. After adding the positional encoding to each peak embedding, they are fed into the encoder layers of the transformer. To prepare the output label during training, the name of the lipid species corresponding to the spectrum is tokenized using regular expressions. Each token is then mapped to its corresponding embedding vector, which is fed into the transformer decoder layer after adding the positional encoding. The final output of the model consists of the name of the predicted lipid species and adduct for each input spectrum. A greedy and a beam search method were implemented as prediction options.

**Figure 1:**
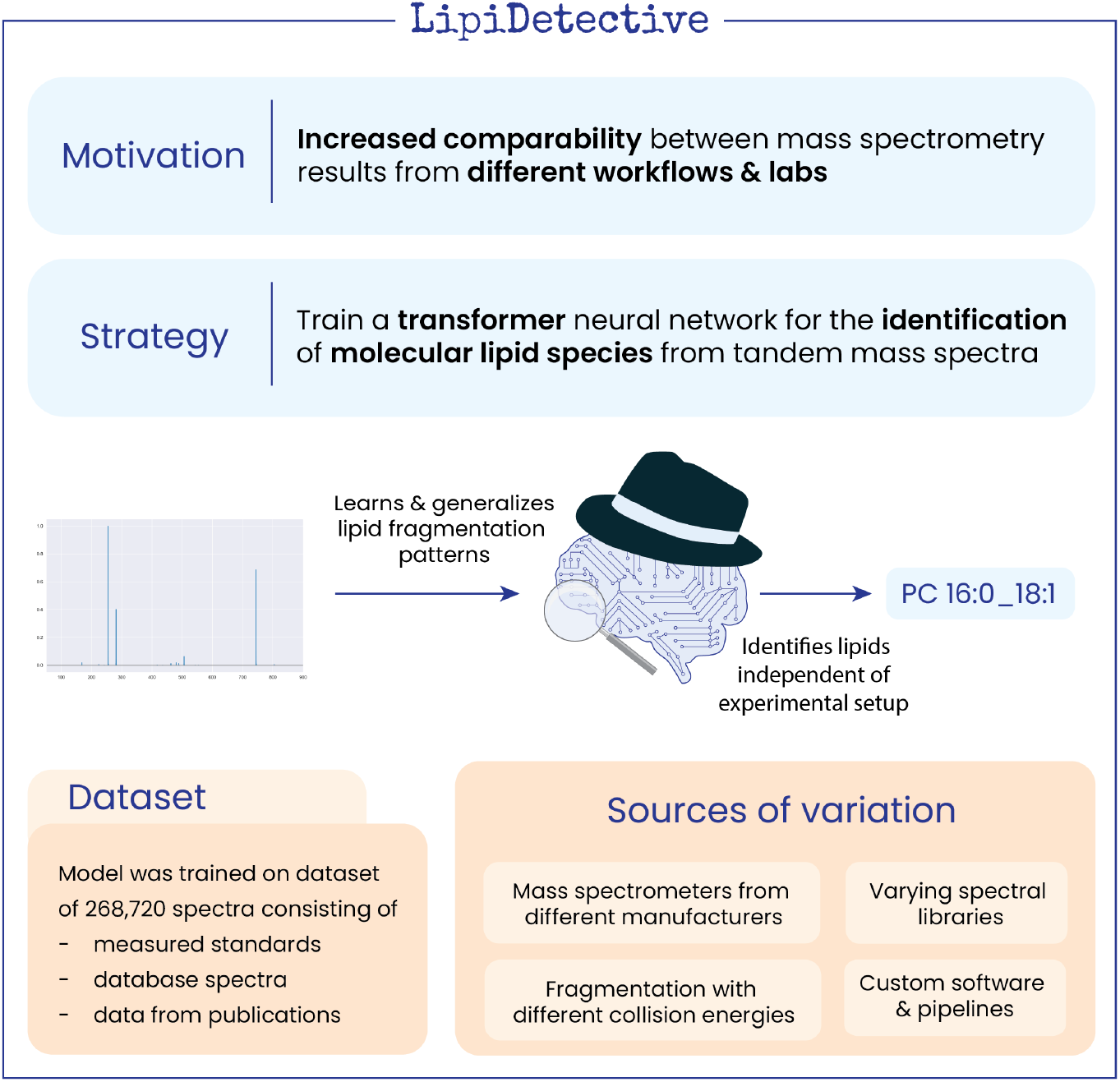
LipiDetective is a deep learning model that can identify molecular lipid species in tandem mass spectra. It was trained on a dataset composed of measured standards, database spectra, and data from publications. As it is able to generalize characteristic lipid fragmentation patterns over varying experimental conditions, it can serve as an additional layer of quality control for identifications and improve comparability between results compared to standard spectral library matching approaches.

**Figure 2:**
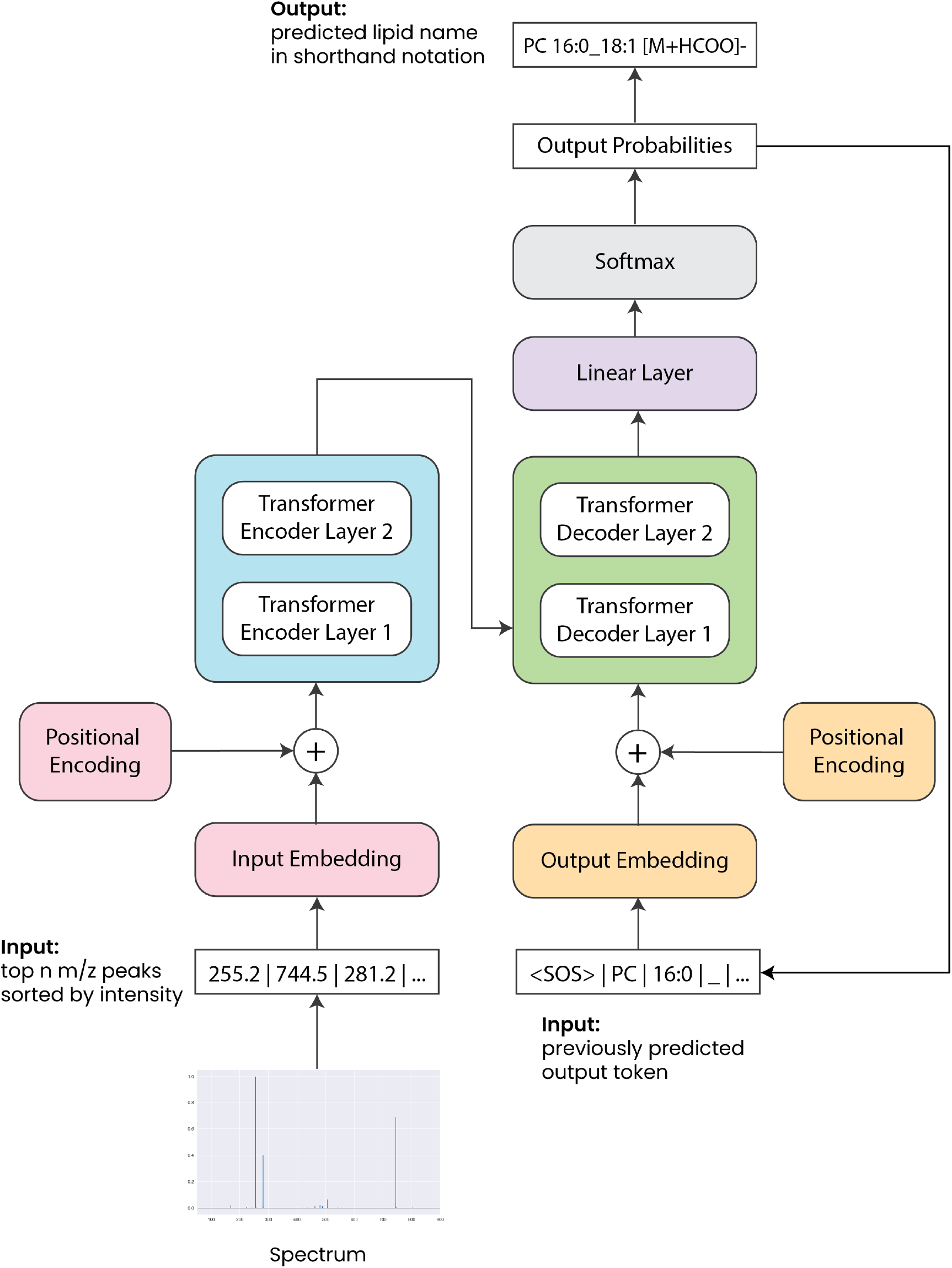
LipiDetective architecture based on the transformer model with an encoder-decoder structure.

### 2.4 Training & Validation

PyTorch Lightning [27] was used for multi-GPU training on a computing cluster. The cluster consists of multiple compute nodes equipped with two or three GPUs. The specific GPU models vary across nodes, including NVIDIA’s Titan Xp, Titan V, and A40. Hyperparameter tuning was performed with Ray Tune [28] to find suitable values for the learning rate, batch size, dropout, embedding size, number of heads and layers, size of the fully connected linear layer, number of input peaks, and decimal point cutoff. All necessary configuration parameters for a training run are stored in a YAML [29] config file, whose save path is the only argument provided to LipiDetective. The different validation splits were also saved as YAML files to provide reproducibility and allow for easy selection. It took around one hour to train the model on the composed dataset with the optimal hyperparameters (Table S3) using early stopping after 15 epochs.

### 2.5 Performance Evaluation

#### 2.5.1 Accuracy Metrics

Two accuracy measurements were implemented to evaluate LipiDetective’s performance. The first one represents total prediction accuracy and compares each predicted token with its corresponding counterpart in the label. Only if all tokens match will the prediction be classified as correct. However, this does not indicate how close the prediction was to the label. For this purpose, the lipid component accuracy was implemented, calculated by dividing the number of correctly predicted tokens by the total tokens. For example, the label `PC 16:0_18:1 [M+HCOO]-`consists of four components: the headgroup `PC`, the first side chain `16:0`, the second side chain `18:1`, and the adduct `[M+HCOO]-`(see Table 1). Suppose the model predicts `PC 16:0_18:0 [M+HCOO]-`, where three out of four components match, it would achieve a lipid component accuracy score of 3/4 = 75%. This provides another useful measure to evaluate how close the model’s predictions are on average to the label.

**Table 1:** Example prediction for determining the lipid component accuracy metric. With three out of four components matching between the label and prediction the lipid component accuracy score is 75 %.

|  |  |  |  |  |  |  |  |  |
| --- | --- | --- | --- | --- | --- | --- | --- | --- |
| **Label** |  | PC |  | 16:0 |  | 18:1 |  | [M+CHOO]- |
| **Prediction** |  | PC |  | 16:0 |  | 18:0 |  | [M+CHOO]- |
| **Match** |  | ✓ |  | ✓ |  | ✗ |  | ✓ |

#### 2.5.2 Interpretability

To understand the impact of individual features on LipiDetective’s predictions, the integrated gradients [30] method was employed as provided by the Captum [31] library. Furthermore, the spectrum embeddings and attention weights generated by the model during inference were extracted and analyzed. Dimensionality reduction using UMAP [32] was performed on the spectrum embeddings for visualization.

## 3 Results and Discussion

### 3.1 Unbalanced lipid class distribution in training data poses a challenge for model evaluation

After preprocessing all MS2 spectra from the four different sources, the final training dataset consists of 268,720 spectra covering 2,306 different molecular lipid species belonging to 31 different lipid classes. Additionally, 2002 MS2 spectra from blank measurements that do not show any target lipid species were added. This was done with the intention that the model would learn to predict nothing if no lipid species was present in a spectrum by directly returning the end-of-sequence token. The distribution of lipid classes in the training data (Supplement Figure S2A) is quite unbalanced, with over 50% of the spectra belonging to phosphatidylcholine (PC), phosphatidylethanolamine (PE), or triacylglycerol (TG) species. Additionally, the composition of lipid classes differs widely between the different sources (Supplement Figure S2B). As the standards dataset consists only of phospholipid measurements, it naturally contributes to their over-representation in the merged dataset. Even before preprocessing, the MITOMICS dataset contains an unusually high percentage of lysophosphatidylethanolamines (LPEs) compared to the other sources. One reason for the variation in lipid composition between the different sources could be that they originate from different matrices, ranging from liver, heart, kidney, conjunctival, and corneal epithelium cells to internal standards and cancer cell lines [33]. Another possibility could be different preanalytical sample handling. An increase of lysophospholipids, including LPEs, can often be observed if unquenched lipid samples are stored for a longer time at room temperature. In general, measurements from a recent study on the NIST plasma composition [7] (Supplement Figure S3) show that an unbalanced distribution of lipid classes for a sample is to be expected with the most prevalently measured classes being PC, PE, and TG.

Even though this data imbalance is expected, it poses a challenge when trying to accurately quantify the model’s performance. DL models generally require large amounts of data to perform well and tend to favor the majority class when using an imbalanced dataset for training. Therefore, the model is expected to perform much better on the highly prevalent lipid classes than on the ones represented by only a few spectra in the training data. This has to be taken into account when trying to evaluate the overall accuracy of its predictions. When splitting the dataset randomly, the model could achieve a high average accuracy if it predicts the abundant lipid classes correctly, even if it completely fails on the rare lipid species, as only a few spectra of the latter would be included in the validation data. For this reason, multiple splits were generated using different strategies (Figure 3A).

**Figure 3:**
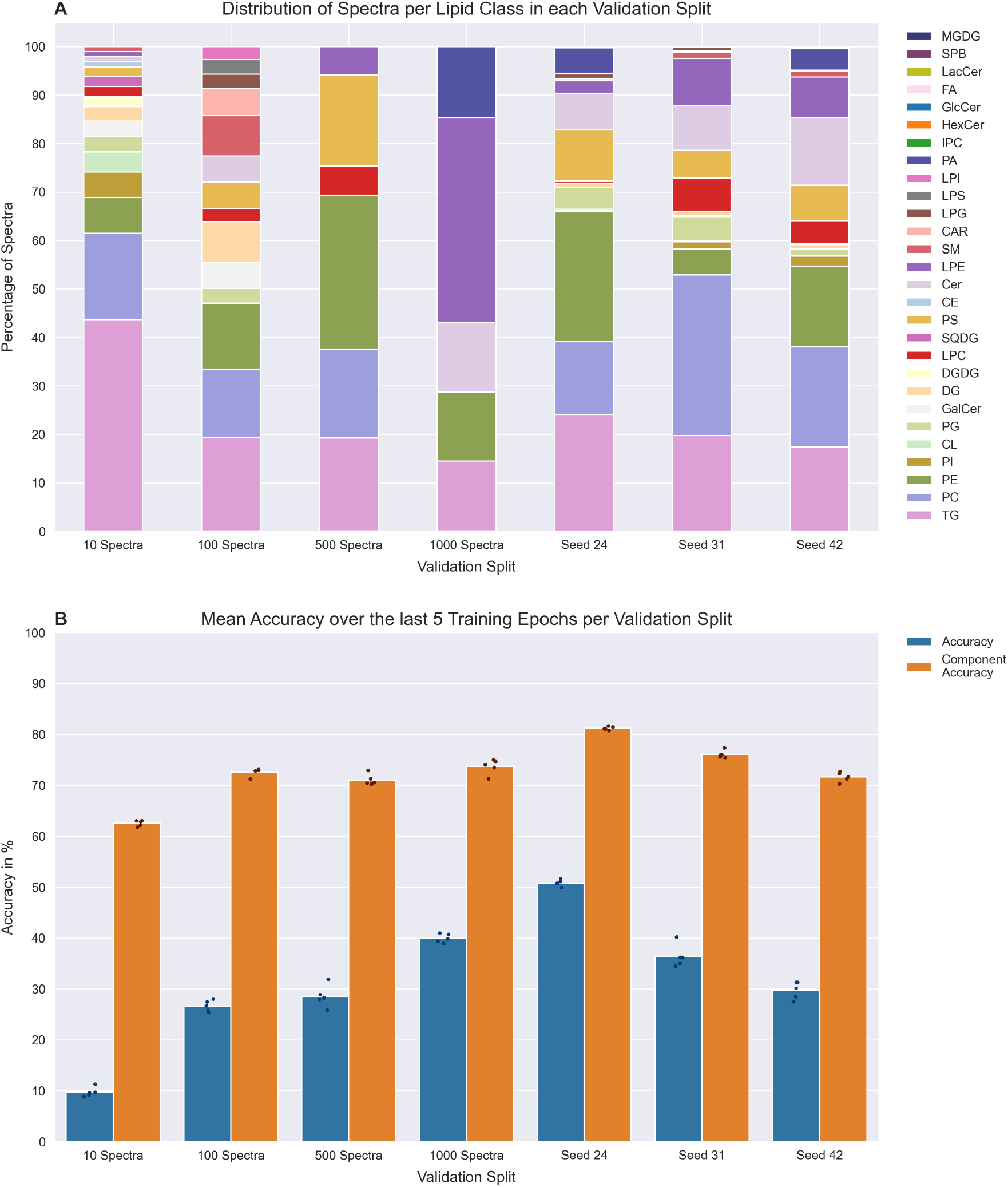
A) Comparison of lipid class distribution in different validation splits. B) Total prediction accuracy and lipid component accuracy of the tuned transformer model for each validation split. Bars represent the mean over the last five epochs, while points indicate the value measured for one epoch.

The general approach for all these splits was to select a set of lipid species, take all their corresponding spectra in the dataset, and exclusively assign them to the validation set. Assuring that the model has never seen any spectra of these species before prevents leakage and allows for rigorous testing of the model’s generalization ability. Three random splits with different random seeds were created using the aforementioned approach to evaluate the influence of class imbalance on performance. An additional four splits were created in which all lipid species with a similar number of spectra (10, 100, 500, and 1000) were assigned to the validation set so that they would influence the validation accuracy roughly the same. Finally, separate splits were created for each lipid class by randomly selecting a number of species for a class to assign to the validation set. This was done for all lipid classes except ethanolaminephosphorylceramides (EPC) and sulfatides (SHexCer), as there were only two spectra each with these classes in the dataset. Comparing the performance on these 36 different splits should allow for a reasonable estimation of the model’s accuracy in general as well as for over- and underrepresented lipid classes.

### 3.2 Performance of LipiDetective varies considerably depending on validation split

Hyperparameter tuning was performed on the 100 spectra validation split as it has a higher lipid class diversity than the 500 and 1000 splits and contains more lipid species with common fatty acid side chain compositions than the 10 split. After determining the optimal hyperparameter settings (Supplement Table S3), new instances of the model were trained from scratch for each of the different validation splits. Accuracy was determined in two ways, as described in the methods section 2.5.1. On average, the model consistently achieves an accuracy of around 87% on the training data. However, the performance varies considerably for the different validation splits.

The total prediction accuracy between the three random splits (Figure 3B) ranged from 29.7% to 50.8%. For the spectra count splits, the accuracy increased with the number of spectra, from the 10 spectra split with the lowest accuracy of 9.7% to the 1000 spectra split with the highest accuracy at 39.9 %. One reason for this might be that the complexity of the validation set decreases with increasing spectra number. The 1000 spectra dataset contains a lower number of different molecular lipid species and, consequently, fewer different lipid classes and fatty acids (Supplement Table S4). Additionally, it makes sense that the lipid species represented with more spectra in the dataset tend to have a more common fatty acid composition. The more often the model has seen a particular fatty acid, the better it should be able to identify it.

Generally, the lipid component accuracy varies less and is much higher than the total prediction accuracy, averaging between 60 % and 80 %. This indicates that even though some predictions might not be completely accurate, the model does not return just some other random lipid but is still able to identify, on average, more than half of the lipid component tokens correctly. This can be confirmed by looking at the actual predictions for the validation dataset. For example, in the last 10 predictions recorded for the validation run with the 100 spectra split the model predicts `DG 18:1_18:2 [M+NH4]+` instead of `DG 18:1_18:1 [M+NH4]+` and `PC 18:1_22:6 [M+H]+` instead of `PC 18:1_22:5 [M+H]+` (Supplement Table S5). These predictions are already very close to the label and differ only by one double bond in one of the fatty acids.

For each individual lipid class validation split, the model achieves the highest total prediction accuracy for PG, followed closely by PE, PC, and phosphatidylserine (PS) (Figure 4). One reason for the excellent performance on PG could be that the training dataset contained only PG spectra measured in negative ion mode. This already removes a relevant source of variation. In addition, there is a much smaller number of only 97 different PG lipid species in the dataset compared to the 428 and 247 different species for PC and PE respectively. However, there seems to be no clear correlation between the number of spectra per lipid species and accuracy. One striking outlier is the model’s exceptionally good performance on diacylglycerol (DG) species, even though the number of DG spectra is relatively low. This indicates that the model is likely transferring a pattern it has learned from a different lipid class onto the DG spectra. Looking at some of the mispredictions in the last training epoch of the run on the DG validation split, one can see that the model often predicts a TG with the same fatty acids as the correct DG label and simply repeats one of the fatty acids (Supplement Table S6). Seeing how similar their spectra are, it is reasonable that the model could learn relevant features for DGs from the TG spectra. The main distinguishing factor between the spectra for these predicted lipids and the correct lipids would be the precursor mass. Increasing the number of DG spectra for the model to train on could help it learn to pay closer attention to the precursor peak to differentiate between TGs and DGs.

**Figure 4:**
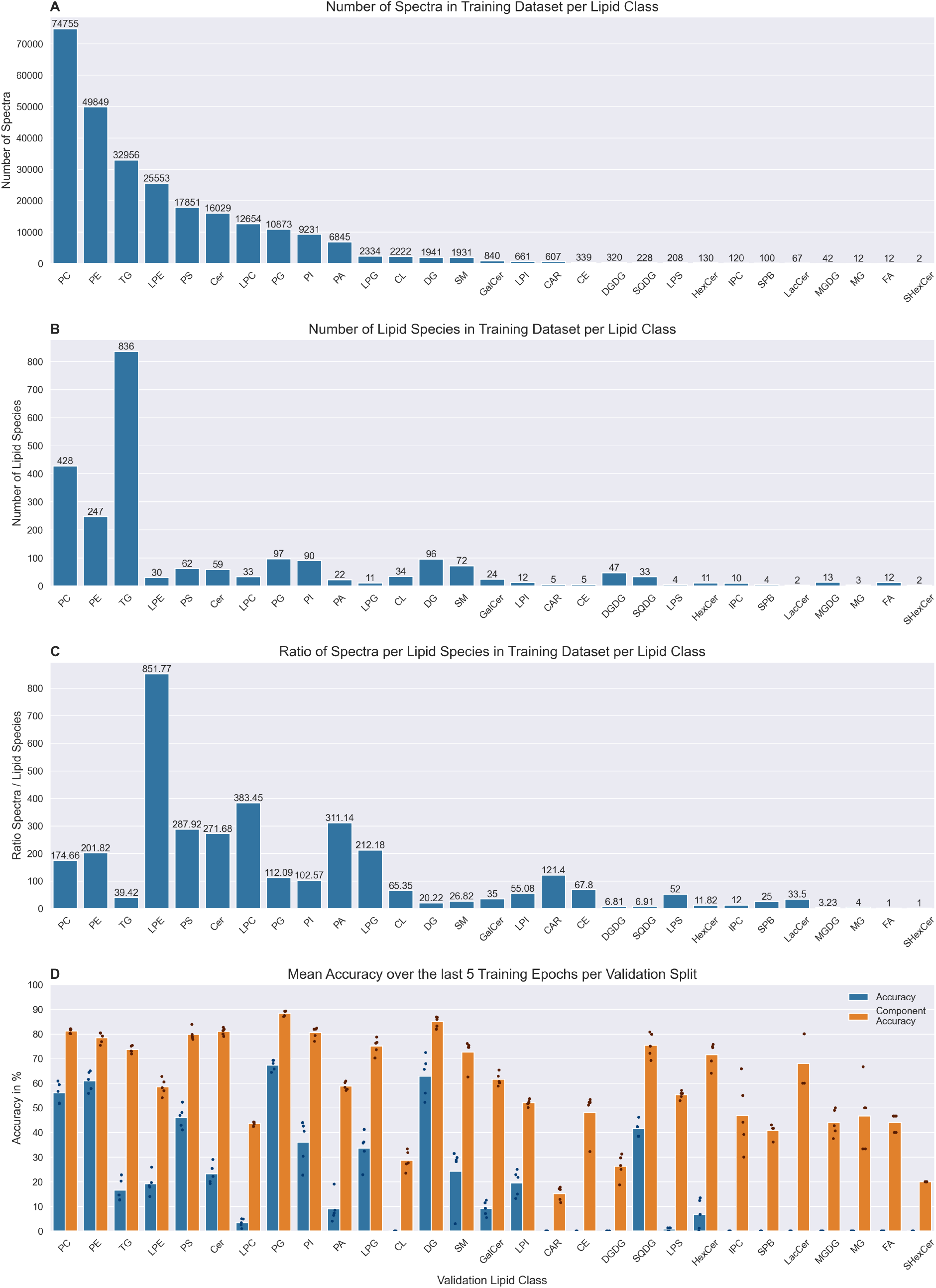
A) Number of spectra per lipid class in the complete training dataset. B) Number of molecular lipid species per lipid class in the training dataset. C) Ratio of spectra/lipid species. D) Performance of LipiDetective on each lipid class validation split.

Overall, it is clear that the lipid class composition of the validation dataset enormously influences the model’s performance. Nevertheless, the presented results are very promising, as the model can achieve high accuracies when identifying lipid species it has never seen before. When using a regular identification pipeline that applies spectrum matching, it would not even be possible to identify an unknown lipid that doesn’t occur in the spectral database. Rule-based methods offer another promising approach to this challenge [34]. However, fragmentation rule sets vary between tools due to differing data sources and instrument dependencies, introducing inconsistencies that hinder comparability. Furthermore, while modern rule-based systems are becoming more capable of adapting to new data, they still rely heavily on established information about lipid fragmentation, potentially overlooking novel informative peaks.

Considering that a relatively small dataset was used for training this neural network, it is very likely that LipiDetective has not yet reached the maximum possible predictive performance. Its best performance on PGs with an accuracy of 67.3 % indicates that there is still a high potential to be harvested by expanding the training dataset to include more spectra of underrepresented lipid species. Another possible influence on the model’s performance is that it was not feasible to manually check the quality of each of the 268,720 spectra and identifications in the training dataset. This could lower the maximal accuracy the model can achieve, and some misidentifications might be due to poor spectrum quality. To improve the quality of the training data, more lipid standard measurements and manually curated spectra could be added.

### 3.3 Encoder spectrum embeddings show that LipiDetective generalizes lipid fragmentation patterns across different experimental conditions

One of the main goals of using a DL model is that the neural network would learn to generalize lipid fragmentation patterns over different experimental conditions. This means the model could identify a lipid correctly, even if it has not previously seen a spectrum of this lipid with that exact collision energy or measured with that specific mass spectrometer. To investigate the model’s abstraction abilities, the spectrum embeddings generated by the encoder were extracted for each lipid in the training dataset and visualized using UMAP (Figure 5).

**Figure 5:**
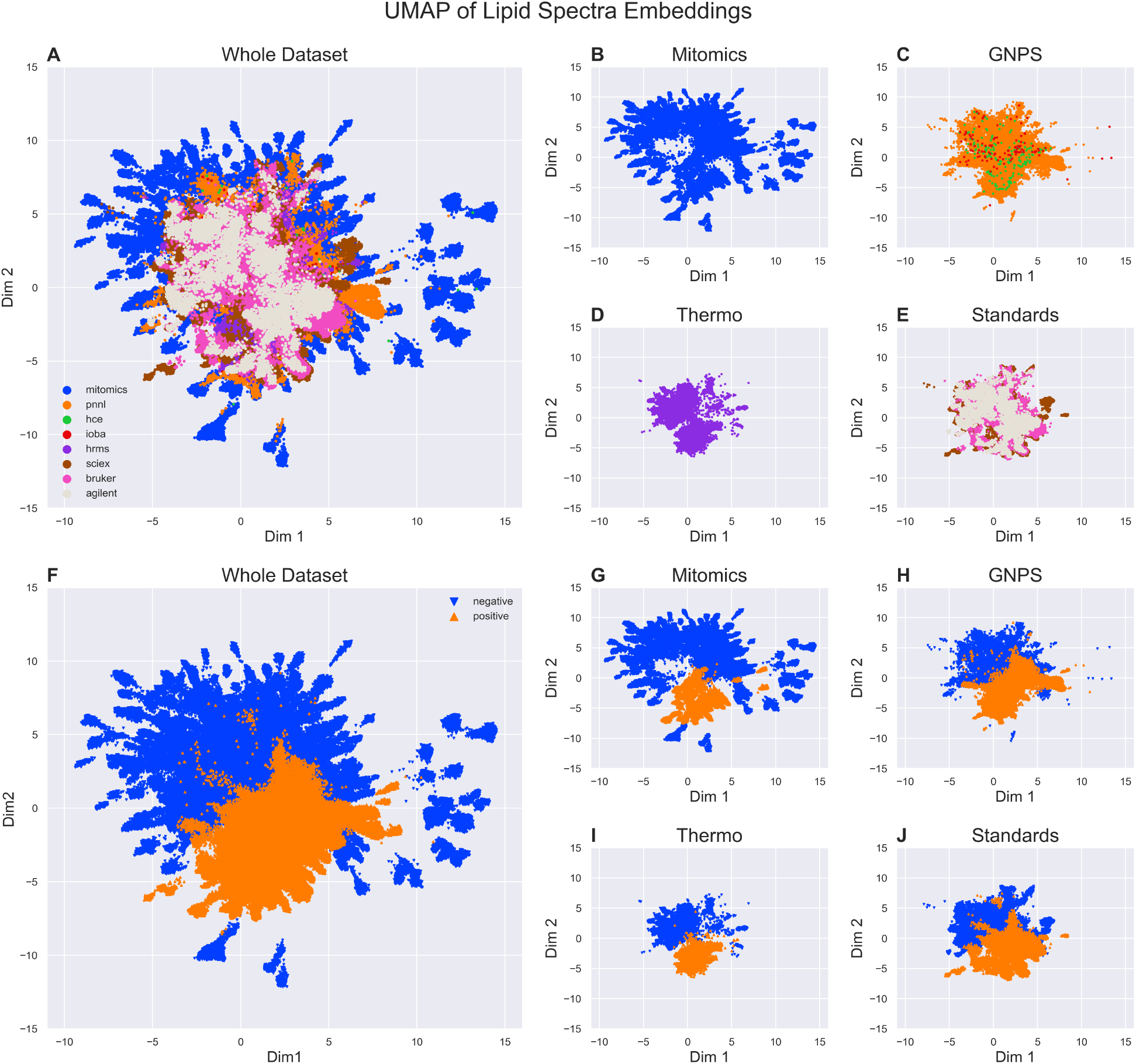
UMAP of spectrum embeddings of A) the whole dataset and separated for each of the sources, B) MITOMICS, C) GNPS, D) Thermo, and E) the phospholipid standards colored by source. The same embeddings were also visualized colored by polarity for F) the whole dataset and each of the sources G) MITOMICS, H) GNPS, I) Thermo and J) the phospholipid standards.

Coloring the UMAP visualization by polarity reveals that the model differentiates between positive and negative measurement mode in the embedding space (Figure 5F). This is unsurprising as the spectra for the same lipid show very different peaks depending on the polarity. However, the embeddings for spectra from different sources overlap remarkably, even though they vary highly in their lipid class composition (Figure 5A). This trend is even more apparent when looking only at the phospholipid standard spectra (Figure 5E), which contain the exact same lipid species measured with mass spectrometers from three different manufacturers, Agilent 6560 IM QTOF-MS, Sciex X500R QTOF-MS, and Bruker maXis UHR-TOF-MS. Although the embeddings of the spectra of the three manufacturers show slight differences and shifts, they mostly overlap. This indicates that the fragmentation patterns between the spectra from different mass spectrometers do not show enough variance to be relevant for the model to differentiate between them.

This is also evident when looking more closely at the embeddings of single species, such as `PC 16:0_18:1`, the most prevalent lipid in the training dataset (Supplement Figure S4). Its embeddings separate into two main large clusters corresponding to positive and negative mode, while the spectra from different sources largely overlap. An exception is one small separate cluster of Sciex spectra measured in the negative mode that is closer to the positive spectra. This is likely due to the different collision energies used when generating the spectra, as all the phospholipid standards were measured with stepped collision energies. Comparing the spectra from each cluster supports the hypothesis that the outlier cluster represents spectra measured at low collision energy, with the precursor peak having the highest intensity and showing barely any fatty acid fragment peaks (Supplement Figure S5). Therefore, it is unsurprising to see them clustering closer together with spectra measured in positive mode, as those also don’t show any fatty acid fragment peaks below 400 m/z.

Overall, the embeddings support the conclusion that LipiDetective achieves one of its primary goals: abstracting the lipid fragmentation patterns independently of the instrument or source. This means that the model does not need to see the spectrum of a lipid under the exact same experimental conditions to identify it correctly. This should allow for an increased comparability between measurements of the same sample by different labs.

### 3.4 Integrated gradients show that Lipidetective considers well-known fragments in its prediction for the corresponding lipid component

To provide more interpretability on the model’s prediction process, the integrated gradients algorithm was used to assign an importance score to each input feature with respect to each output token. Figure 6 shows these importance scores for a spectrum of `PC 16:0_18:1` measured in negative mode with the adduct `[M+HCOO]-`. The y-axis displays the input m/z values, which are annotated with expected fragments for this lipid species taken from Alex^123^. The x-axis indicates the predicted output tokens. Interestingly, the PC headgroup loss at 744.5 m/z as well as the precursor mass at 804.5 m/z achieve high importance scores for the prediction of the PC token. A human observer would also use these peaks as an initial indication of a PC in negative mode, supporting the hypothesis that the model evaluates the spectra in a manner similar to a human. Additionally, the model also considers isotopic peaks such as 746.5 m/z for the PC headgroup prediction. The peak at 255.2 m/z highly influences the prediction of `16:0` as the first fatty acid, which makes sense as the peak corresponds to the `16:0` fatty acid fragment. Similarly, the input peak at 281.2 m/z corresponding to the known fatty acid `18:1` fragment shows the highest importance score for the prediction of `18:1` as the second side chain. This indicates that LipiDetective has, in fact, captured the pattern that certain peaks in a spectrum are characteristic of fragments that correspond to particular components of a lipid. There are some other peaks, such as 224.0 m/z, that are also relevant for the prediction but for which there is no annotation in the Alex^123^ database. Understanding which molecule they might correspond to could reveal new information about the fragmentation processes of lipids and help create better reference spectra. In fact, literature research reveals that this peak most likely corresponds to a fragment comprising the PC headgroup and glycerol backbone [35]. All this shows that the model seems to primarily focus on human interpretable peaks that can be directly correlated to specific lipid fragments and corresponding isotopic peaks.

**Figure 6:**
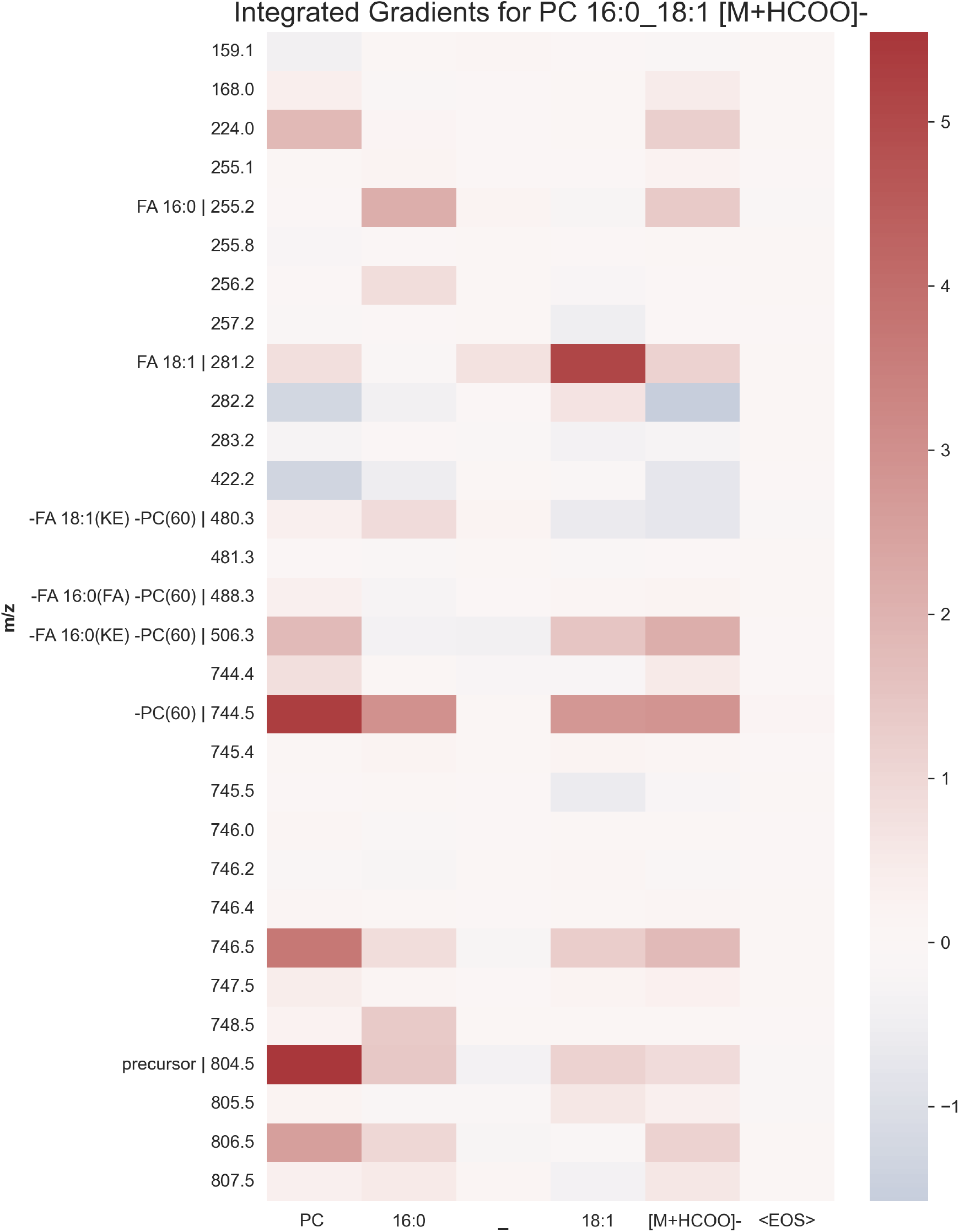
Integrated Gradients for a spectrum showing PC 16:0_18:1 [M+HCOO]-. Important peaks corresponding to known fragments in the Alex^123^ database are annotated.

These importance scores could also be used to extract the most important peaks over all spectra the model trained on and create an in silico spectrum from them. For this purpose, the importance scores were calculated for each spectrum of a selected lipid species with a specific adduct. Then, they were summed over all output tokens and all spectra to create a single importance score for each input feature. The highest importance score was then set to one and used to scale the other scores accordingly. Finally, all input features that achieved less than 1 % of the maximum score were filtered out. Visualizing the remaining m/z values with the scaled importance scores as intensities results in a plot that looks incredibly similar to a mass spectrum with a medium-level collision energy (Supplement Figure S7). This is surprising as the model never sees the actual values of the peak intensities. The only indication it has of the intensities is by ordering the input vector with the m/z values by descending intensity. The in silico spectrum for `PC 16:0_18:1 [M+HCOO]-`generated using this method contains 24 m/z values, whereas most predicted spectra typically show fewer peaks. Additionally, it indicates the relevance of the peaks by providing the importance values as intensities.

Interesting information might also be gained from using integrated gradients on a misprediction. A spectrum of `TG 22:5_22:6_22:6 [M+NH4]+` from PNNL was chosen to showcase such an example. The prediction of the LipiDetective model for this particular spectrum was `TG 16:0_22:6_22:6 [M+NH4]+`, so it mispredicted the first fatty acid as `16:0` instead of `22:5`. Looking at the integrated gradients (Supplement Figure S8) one can see that this prediction was mainly influenced by the peak at 313.2 m/z. When cross-referencing with the Alex^123^ database, this peak corresponds precisely to the `22:5` fatty acid fragment. At first glance, it seems surprising that the model would choose this as an informative peak to predict `16:0`. However, when looking at the expected fragments for `TG 16:0_22:6_22:6 [M+NH4]+` in the Alex^123^ database, the exact same peak appears at 313.2 m/z for the fragment `FA 16:0(+C3H6O2)`. These peaks would be differentiable by the second decimal place, as they correspond to 313.25 m/z for `FA 22:5` and 313.27 m/z for `FA 16:0(+C3H6O2)`. However, LipiDetective currently only uses m/z values with a precision of two decimal points. A simple solution for this issue would then be to increase the number of decimal places for the m/z values fed into the model. Unfortunately, this notably decreased performance on the validation splits. One likely reason for this is the relatively small dataset size, resulting in the model’s inability to adjust to the enormous increase of possible input features from 16,000 to 160,000 possible peaks. Additionally, when increasing the decimal place, each peak appears less frequently, and the learned peak embeddings are likely not quite as informative.

It is also surprising that the model mispredicted the fatty acid position as it does seem to consider the precursor mass at 1024.1 m/z. This is likely due to the imbalanced dataset, which leads to a bias towards the more commonly occurring fatty acids. For TGs in the training data, the fatty acid `16:0` occurs 12,794 times in the first position compared to `22:5`, which occurs only two times. When disregarding the position, `16:0` occurred 21,131 times compared to only 583 times for `22:5`. This means the model is much more likely to predict a more common fatty acid such as `16:0` if it can find reasonably fitting peaks. This could also be remedied to a certain extent by expanding the dataset to cover more uncommon fatty acids.

Using integrated gradients showed unwanted biases in the dataset that affect the prediction. This needs to be considered when using LipiDetective to identify more uncommon lipids. To give the user a better indication of the confidence of the prediction, a function was implemented so that LipiDetective returns the top three predictions for a spectrum, including an overall probability score between zero and one for each prediction. For example, for the misprediction `TG 16:0_22:6_22:6 [M+NH4]+`, the model’s probability score reached only around 0.4747, whereas it returns a score of 0.9970 for the correct prediction of the `PC 16:0_18:1 [M+HCOO]-`spectra in Figure 6. This feature can serve as an additional quality check of the model’s prediction, allowing the user to determine their own confidence threshold.

### 3.5 Visualization of attention weight matrices gives further insight into LipiDetective’s input processing

As the LipiDetective model is based on the transformer architecture, it utilizes attention, which can be used as an additional tool to improve its interpretability. High attention weights can hint at key inputs that provide contextual information for other peaks or even the whole spectrum. Figure 7 visualizes the attention for all four attention heads in both encoder layers for `PC 16:0_18:1` measured in negative mode with adduct `[M+HCOO]-`at a collision energy of 20.0 eV.

**Figure 7:**
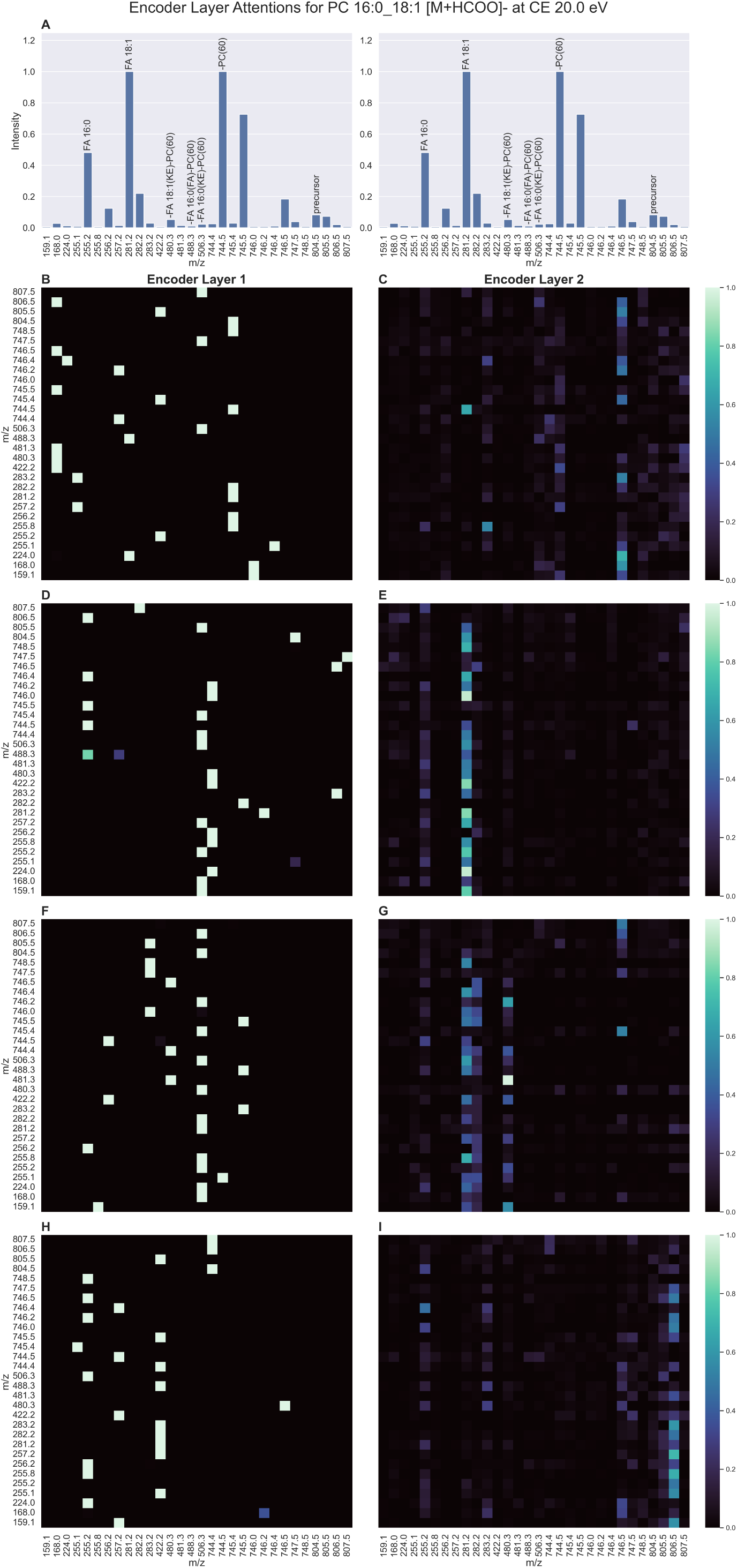
A) Spectrum of PC 16:0_18:1 [M+HCOO]-at a collision energy of 20 eV. B) Corresponding matrices of attention weights, with each row representing a different attention head and column 1 corresponding to the first and column 2 to the second encoder layer.

In the first encoder layer, most input peaks exhibit one high attention value focused on one other input peak. However, the attention weights in the second encoder layer are much more diffusely distributed across the input peaks. This could mean that the model focuses primarily on capturing local dependencies within the input data in the first encoder layer. Specific peaks exhibiting high attention to each other may indicate strong local correlations in the mass spectrum. Then, in the subsequent layer, the model integrates these local features to build a more global representation of the input, leading to a more diffuse attention pattern.

Making sense of the local dependencies between peaks is quite challenging. However, it is clear that each attention head of the second encoder layer seems to focus on a different aspect of the input spectrum. The first attention head focuses mainly on the headgroup, while the second concentrates on the fatty acid peaks. The third attention head also pays attention to the fatty acid peaks while also including the fragments representing the combined fatty acid and headgroup loss. The fourth attention head mainly focuses on the precursor mass. The peaks exhibiting high attention values in the second encoder layer show nearly none in the first encoder layer and vice versa. All this supports the theory that LipiDetective recognizes representative fragment peaks and then uses them in the second encoder layer to provide overall context for the other peaks in the mass spectrum.

## 4 Conclusion

To our knowledge, LipiDetective is the first attempt at leveraging the capabilities of DL to identify molecular lipid species from tandem mass spectra. The approach that comes closest to using deep neural networks for lipid identification is MSNovelist, which combines fingerprint prediction with an encoder–decoder neural network to generate structures de novo from tandem mass spectra [36]. However, this tool is targeted mostly towards metabolomics, which is evident by its limitation to molecules below 1000 Da, thereby excluding many TG species and cardiolipins. Additionally, as its main goal is the creation of structures, it does not deliver lipid identifications in standardized lipid shorthand nomenclature. This positions LipiDetective as a unique and specialized tool for lipidomics, enhancing the utility and accessibility of lipidomics data by delivering molecular lipid species level identifications in a standardized format. Through comprehensive evaluation, it was demonstrated that LipiDetective recognizes representative fragment peaks and can abstract the lipid fragmentation patterns independently of the instrument or laboratory. This allows LipiDetective to identify lipid species for which it has never even seen a spectrum before.

Naturally, the novelty of this approach comes along with limitations. It was shown that biases in the dataset affect the predictions and that LipiDetective prefers to predict more common fatty acids prevalent in the training data. Considering that the training dataset for LipiDetective was relatively small, it can be assumed that the model has not yet reached its highest performance level. These issues will be addressed by improving the quality of the training data by adding more lipid standard measurements and manually verified spectra. As this is a time-intensive endeavor, more immediate assistance is needed to help the user evaluate the model’s confidence in a prediction. For this purpose, LipiDetective provides a probability score between zero and one for each identified lipid.

One planned improvement for the model is the integration of additional constraints during prediction. The beam search prediction process can be guided to ensure the predicted tokens generate a lipid species that always matches the precursor mass of the spectrum. This could be achieved by associating a mass equivalent with each token and maintaining a running sum of the total token mass for each beam throughout the prediction process. If the difference between the precursor mass and the calculated mass for a prediction surpasses a threshold value, the prediction would be discarded. This could eventually be combined with user-specified retention time ranges for certain lipid classes, so that only predictions with matching class tokens for the defined retention times are retained.

This first-of-its-kind DL model represents an opportunity for researchers to build upon. The pre-trained LipiDetective model can easily be retrained on additional data, including specific lipid species or more uncommon lipid classes, to improve its performance for targets of interest. LipiDetective is also not limited to the lipid classes or adducts it has trained for. Further lipid class or adduct tokens can easily be added to the vocabulary file that LipiDetective uses. Updated models can then be saved and uploaded with the experimental data of a publication to provide reproducibility, allowing other laboratories to take advantage of such specialized models. Ideally, the training data would also be provided so that it will be possible to develop a model that was trained on a large corpus of spectra from many different laboratories using varying experimental setups.

Another advantage of LipiDetective is its ability to perform predictions quickly. While the exact speed depends on the hardware, LipiDetective can typically annotate files containing around a hundred spectra in less than five seconds. With only around 616,000 parameters, the saved models are highly compact, occupying approximately 2.5 MB of memory. This low-latency prediction and minimal memory footprint make LipiDetective suitable for deployment directly on MS or attached computing devices. Real-time identification of compounds could be particularly beneficial in clinical diagnostics and other fields where immediate results are critical.

In conclusion, this study showcases the potential of DL models to advance lipidomics research by providing increased comparability between measurements by different laboratories. LipiDetective represents a significant first step in this direction by providing a foundation on which the lipidomics community can build.

## Data Availability Statement

All previously unpublished data for this project will be made publicly available on MetaboLights.

## Code Availability Statement

The LipiDetective source code can be accessed on GitHub via https://github.com/LipiTUM/lipidetective.

## Author Contributions

JKP supervised the project and secured the funding. MW supplied the phospholipid standard dataset. LH and RG provided the identified data for the Thermo dataset. VW, NK, and JKP planned and conceptualized the work. FM implemented a transformer prototype. VW designed and implemented the framework and models. VW ran all evaluations. VW and JKP wrote the manuscript. All authors read, reviewed, and accepted the manuscript in its final form.

## Acknowledgments

This project was funded by the Bavarian State Ministry of Science and the Arts in the framework of the Bavarian Research Institute for Digital Transformation (bidt; JKP, VW, NK, FM: Junior Research Group LipiTUM) and the Deutsche Forschungsgemeinschaft (DFG, German Research Foundation) – 422216132.

## List of Abbreviations

ANN: artificial neural network
CID: collision-induced dissociation
DDA: data-dependent aquisition
DG: diacylglycerol
DIA: data-independet acquisition
DL: deep learning
EPC: ethanolaminephosphorylceramides
ESI: electrospray ionization
GNPS: Global Natural Products Social Molecular Networking
HCD: high-energy collisional dissociation
HDF5: Hierarchical Data Format 5
LMSD: Lipid Maps Structure Database
LPE: lysophosphatidylethanolamine
MITOMICS: mitochondrial orphan protein multi-omic CRISPR screen
MS: mass spectrometry
MS2: tandem mass spectrometry
m/z: mass-to-charge
MRM: multiple reaction monitoring
PC: phosphatidylcholine
PE: phosphatidylethanolamine
PG: phosphatidylglycerol
PNNL: Pacific Northwest National Lab
PS: phosphatidylserine
SHexCer: sulfatides
SRM: standard reference material
TG: triacylglycerol

