## Supplements for "LipiDetective a deep learning model for the identification of molecular lipid species in tandem mass spectra"

### Supplement

***Table S2:*** *Lipid species that were measured to create the phospholipid standards data source.*

| Nr | Lipid Species | Positive Adducts | Negative Adducts |
|---|---|---|---|
| 1 | PA 14:0_14:0 | [M+Na]+ | [M-H]- |
| 2 | PA 16:0_16:0 | [M+Na]+ | [M-H]- |
| 3 | PA 16:0_18:1 | [M+Na]+ | [M-H]- |
| 4 | PA 16:0_18:2 | [M+Na]+ | [M-H]- |
| 5 | PA 16:0_20:4 | [M+Na]+ | [M-H]- |
| 6 | PA 16:0_22:6 | [M+Na]+ | [M-H]- |
| 7 | PA 17:0_17:0 | [M+Na]+ | [M-H]- |
| 8 | PA 18:0_18:0 | [M+Na]+ | [M-H]- |
| 9 | PC 14:0_14:0 | [M+H]+ | [M+HCOO]- |
| 10 | PC 15:0_15:0 | [M+H]+ | [M+HCOO]- |
| 11 | PC 16:0_16:0 | [M+H]+ | [M+HCOO]- |
| 12 | PC 16:0_18:0 | [M+H]+ | [M+HCOO]- |
| 13 | PC 16:0_18:1 | [M+H]+ | [M+HCOO]- |
| 14 | PC 16:0_18:2 | [M+H]+ | [M+HCOO]- |
| 15 | PC 16:0_20:4 | [M+H]+ | [M+HCOO]- |
| 16 | PC 16:0_22:6 | [M+H]+ | [M+HCOO]- |
| 17 | PC 17:0_17:0 | [M+H]+ | [M+HCOO]- |
| 18 | PC 18:0_18:0 | [M+H]+ | [M+HCOO]- |
| 19 | PC 18:0_18:1 | [M+H]+ | [M+HCOO]- |
| 20 | PC 18:0_18:2 | [M+H]+ | [M+HCOO]- |
| 21 | PC 18:0_20:4 | [M+H]+ | [M+HCOO]- |
| 22 | PC 18:0_22:6 | [M+H]+ | [M+HCOO]- |
| 23 | PC 20:0_20:0 | [M+H]+ | [M+HCOO]- |
| 24 | PC 22:0_22:0 | [M+H]+ | [M+HCOO]- |
| 25 | PE 14:0_14:0 | [M+H]+ | [M-H]- |
| 26 | PE 15:0_15:0 | [M+H]+ | [M-H]- |
| 27 | PE 16:0_16:0 | [M+H]+ | [M-H]- |
| 28 | PE 16:0_18:1 | [M+H]+ | [M-H]- |
| 29 | PE 16:0_18:2 | [M+H]+ | [M-H]- |
| 30 | PE 16:0_20:4 | [M+H]+ | [M-H]- |
| 31 | PE 16:0_22:6 | [M+H]+ | [M-H]- |
| 32 | PE 17:0_17:0 | [M+H]+ | [M-H]- |
| 33 | PE 18:0_18:0 | [M+H]+ | [M-H]- |
| 34 | PE 18:0_18:1 | [M+H]+ | [M-H]- |
| 35 | PE 18:0_18:2 | [M+H]+ | [M-H]- |
| 36 | PE 18:0_20:4 | [M+H]+ | [M-H]- |
| 37 | PE 18:0_22:6 | [M+H]+ | [M-H]- |
| 38 | PG 14:0_14:0 | | [M-H]- |
| 39 | PG 15:0_15:0 | | [M-H]- |
| 40 | PG 16:0_16:0 | | [M-H]- |
| 41 | PG 16:0_18:1 | | [M-H]- |
| 42 | PG 16:0_18:2 | | [M-H]- |
| 43 | PG 16:0_20:4 | | [M-H]- |
| 44 | PG 16:0_22:6 | | [M-H]- |
| 45 | PG 17:0_17:0 | | [M-H]- |
| 46 | PG 18:0_18:0 | | [M-H]- |
| 47 | PS 14:0_14:0 | [M+H]+ | [M-H]- |
| 48 | PS 16:0_16:0 | [M+H]+ | [M-H]- |
| 49 | PS 16:0_18:1 | [M+H]+ | [M-H]- |
| 50 | PS 16:0_18:2 | [M+H]+ | [M-H]- |
| 51 | PS 16:0_20:4 | [M+H]+ | [M-H]- |
| 52 | PS 16:0_22:6 | [M+H]+ | [M-H]- |
| 53 | PS 17:0_17:0 | [M+H]+ | [M-H]- |
| 54 | PS 18:0_18:0 | [M+H]+ | [M-H]- |

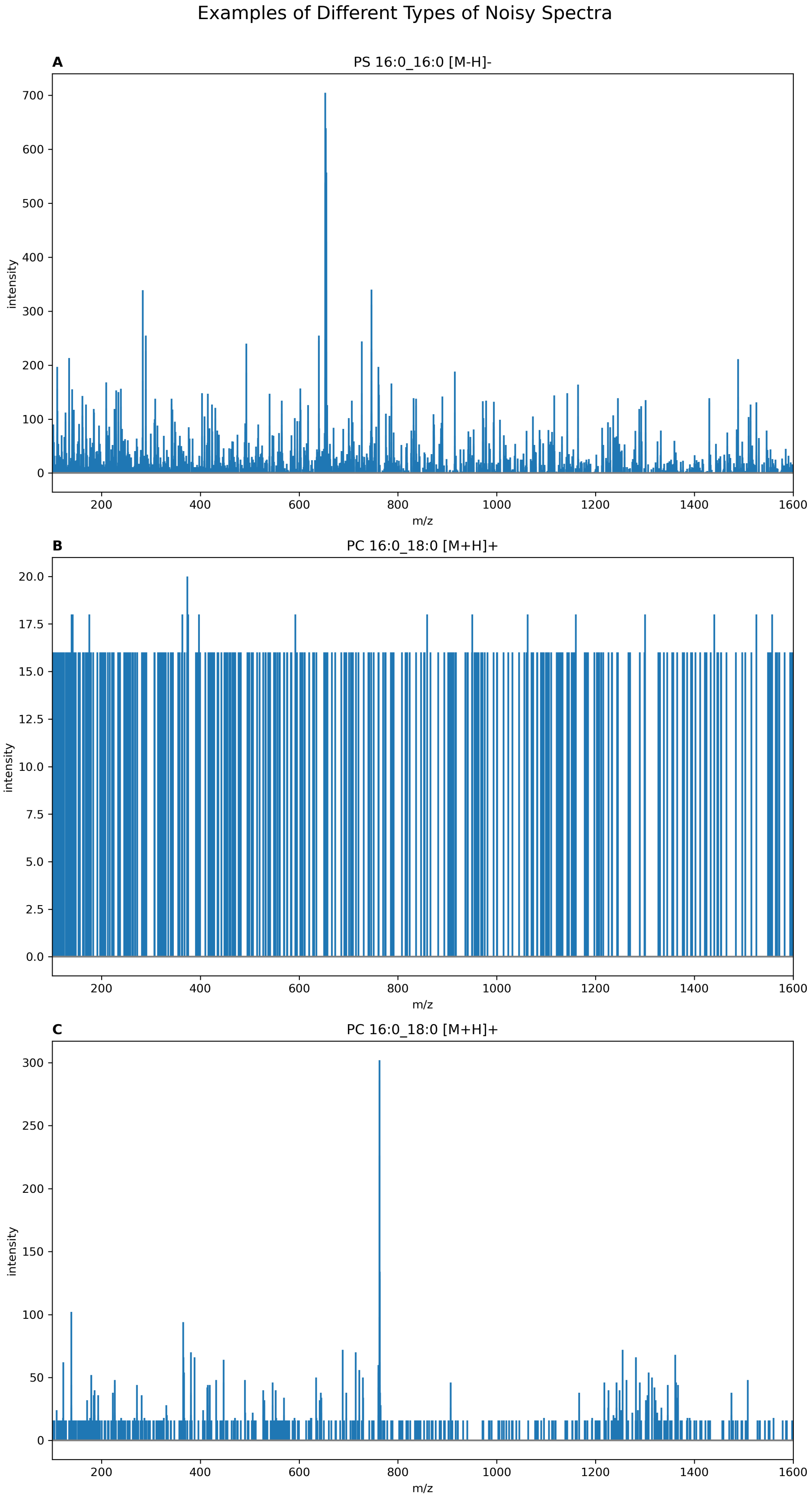


***Figure S1:*** *Examples of noisy spectra filtered from the training dataset using the base peak normalized median intensity. A) MS2 spectrum not containing the target lipid due to an accidentally selected precursor mass of 743.5 Da instead of the true precursor mass of 734.5 Da for lipid standard PS 16:0_16:0 [M-H]- (0.022 normalized median intensity). B) MS2 spectrum of standard PC 16:0_18:1 with adduct [M+H]+ and selected precursor mass of 760.6 Da that seems to show only noise (0.800 normalized median intensity). C) A spectrum from the same file as B) with a very low signal-to-noise ratio showing some target compound fragment peaks (0.020 normalized median intensity).*

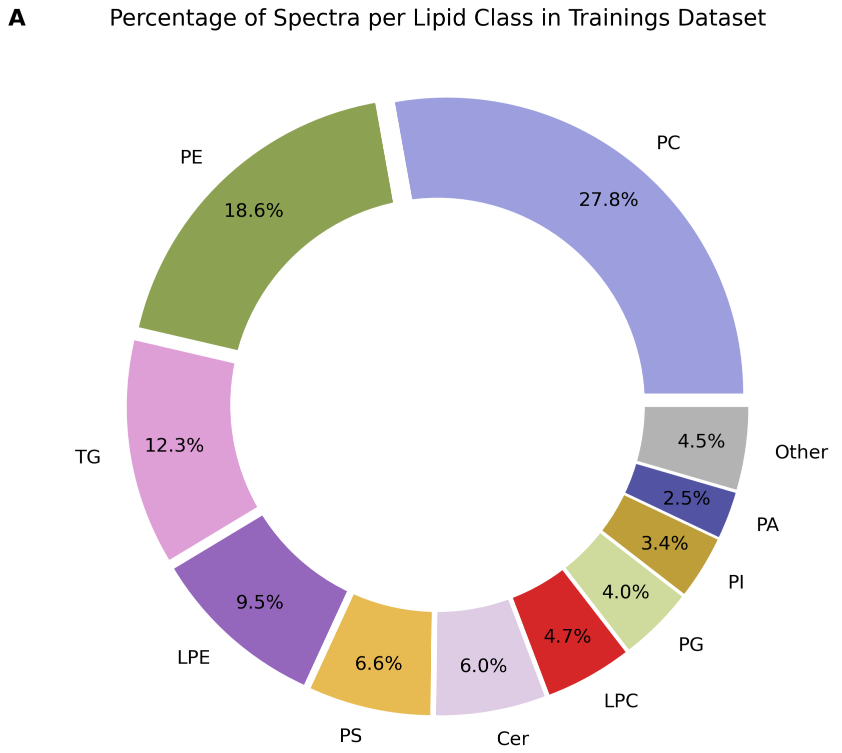


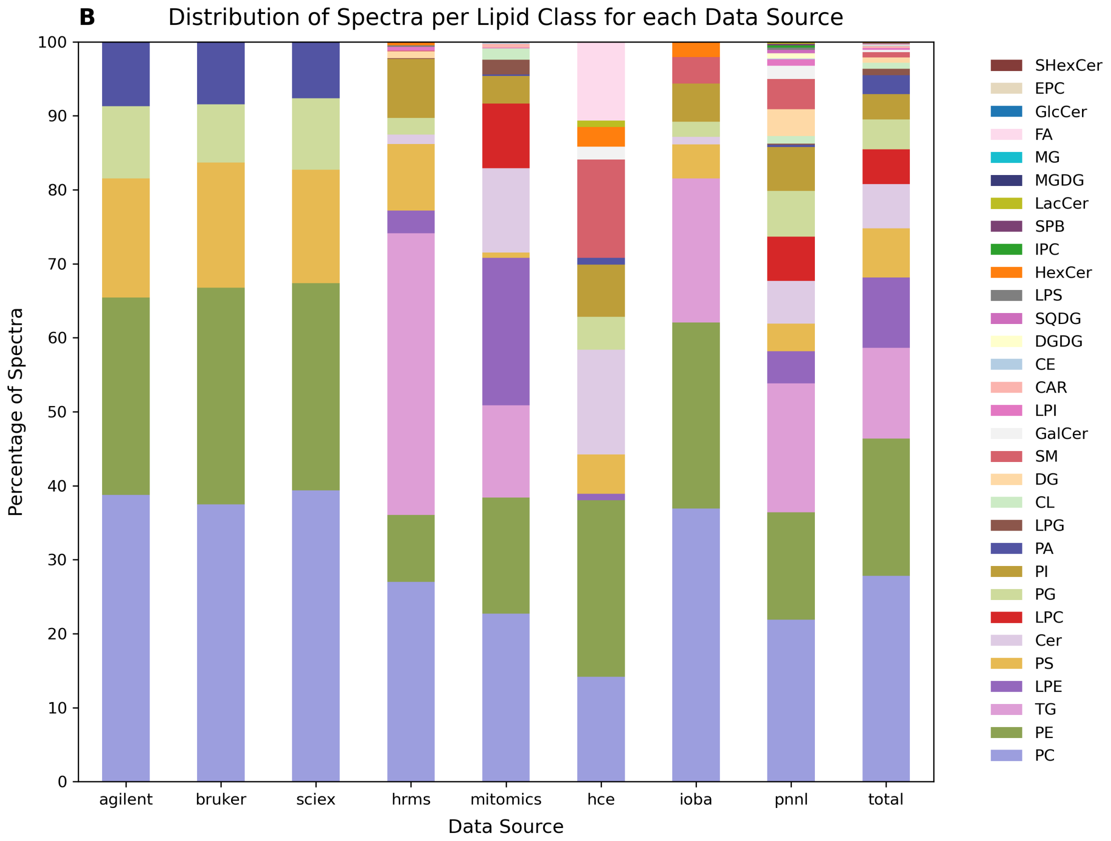


***Figure S2:*** *A) Lipid class distribution of spectra in the training dataset. B) Distribution of lipid classes for each source. Variation between the Agilent, Sciex, and Bruker sources is due to the noisy spectra filtering as described in methods section 2.2.2.*

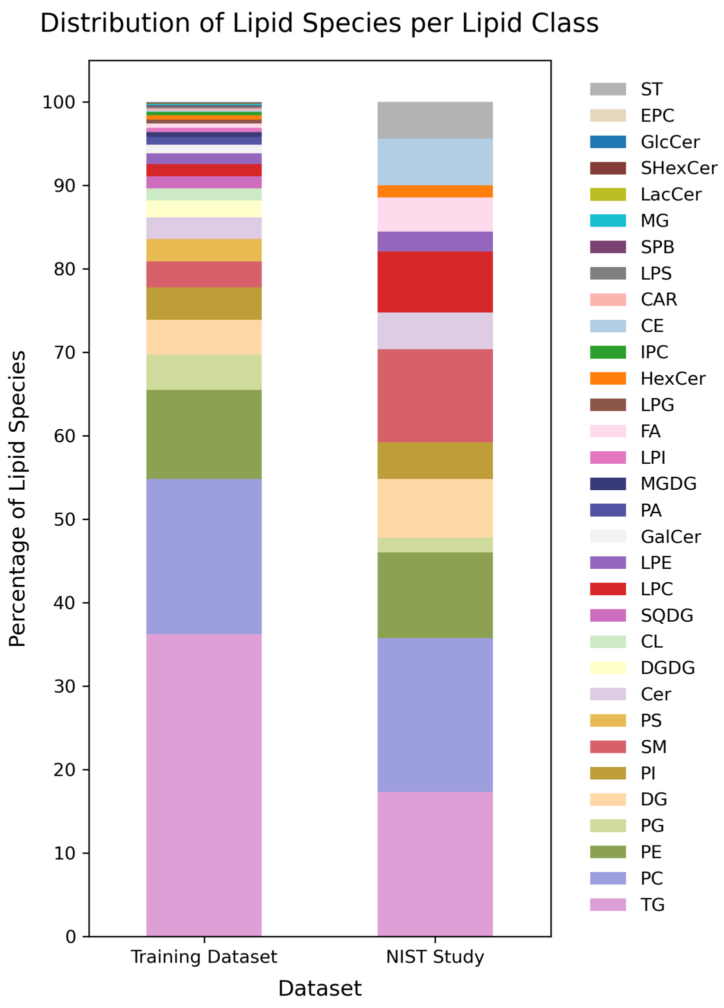


***Figure S3:*** *Comparison of lipid class distribution in training data vs NIST plasma identifications.*

***Table S3:*** *Hyperparameter values chosen after tuning and used for all presented runs.*

| Hyperparameter | Optimum |
|---|---|
| Learning rate | 0.004 |
| Learning rate step | 2 |
| Dropout | 0.1 |
| Batch size | 512 |
| Epochs | 15 |
| Number of peaks | 30 |
| Decimal places | 1 |
| Embedding dimension | 32 |
| Number of heads | 4 |
| Number of layers | 2 |
| Dimension of Feedforward Network | 256 |

***Table S4:*** *Number of different lipid species, lipid classes, and fatty acids in each validation split by spectra count.*

| Val Split | Nr Lipid Species | Nr Lipid Classes | Nr Fatty Acids |
|---|---|---|---|
| 10 Spectra | 93 | 16 | 42 |
| 100 Spectra | 36 | 14 | 22 |
| 500 Spectra | 16 | 6 | 15 |
| 1000 Spectra | 7 | 5 | 6 |

***Table S5:*** *Last 10 predictions recorded for the validation run with the 100 spectra split.*

| Nr | Prediction | Label | Component Accuracy | Correct |
|---|---|---|---|---|
| 1 | TG 14:0_14:0_16:0 [M+NH4]+ | TG 14:0_16:0_16:0 [M+NH4]+ | 0.8 | FALSE |
| 2 | PE P-18:0_20:3 [M+H]+ | PE P-18:0_20:3 [M+H]+ | 1.0 | TRUE |
| 3 | PC 18:1_22:6 [M+H]+ | PC 18:1_22:5 [M+H]+ | 0.75 | FALSE |
| 4 | PE O-18:0_18:1 [M-H]- | PE 17:1_18:1 [M-H]- | 0.75 | FALSE |
| 5 | TG 18:1_18:1_20:3 [M+NH4]+ | TG 16:0_18:1_22:4 [M+NH4]+ | 0.6 | FALSE |
| 6 | PG 18:1_20:4 [M-H]- | PG 18:1_20:4 [M-H]- | 1.0 | TRUE |
| 7 | DG 18:1_18:1 [M+NH4]+ | DG 18:1_18:1 [M+NH4]+ | 1.0 | TRUE |
| 8 | PE P-18:0_18:2 [M-H]- | PE 17:1_18:1 [M-H]- | 0.5 | FALSE |
| 9 | PG 18:1_20:4 [M-H]- | PG 18:1_20:4 [M-H]- | 1.0 | TRUE |
| 10 | DG 18:1_18:2 [M+NH4]+ | DG 18:1_18:1 [M+NH4]+ | 0.75 | FALSE |

***Table S6:*** *Last 10 mispredictions recorded for the validation run with the DG lipid class split.*

| Nr | Prediction | Label | Component Accuracy |
|---|---|---|---|
| 1 | TG 18:3_18:3_21:0 [M+NH4]+ | DG 18:2_18:4 [M+NH4]+ | 0.25 |
| 2 | TG 18:1_18:2_22:0 [M+NH4]+ | DG 18:1_18:2 [M+NH4]+ | 0.75 |
| 3 | TG 20:5_20:5_20:5 [M+NH4]+ | DG 20:5_20:5 [M+NH4]+ | 0.75 |
| 4 | TG 18:1_18:2_18:2 [M+NH4]+ | DG 18:1_18:2 [M+NH4]+ | 0.75 |
| 5 | TG 18:1_18:2_18:2 [M+NH4]+ | DG 18:1_18:2 [M+NH4]+ | 0.75 |
| 6 | TG 18:1_18:2_19:1 [M+NH4]+ | DG 18:1_18:2 [M+NH4]+ | 0.75 |
| 7 | TG 8:0_18:1_28:2 [M+NH4]+ | DG 18:1_18:2 [M+NH4]+ | 0.5 |
| 8 | DGDG 18:1_18:2 [M+NH4]+ | DG 18:1_18:2 [M+NH4]+ | 0.75 |
| 9 | LPE 16:0 [M-H]- | DG 18:1_18:2 [M+NH4]+ | 0.0 |
| 10 | Cer 18:0;O2/16:0 | DG 18:0_18:1 [M+NH4]+ | 0.25 |

***Table S7:*** *Number of lipid species and spectra for each data source as well as average number of peaks per spectrum per source.*

| Source | Nr Lipid Species | Nr Spectra | Mean Nr Peaks |
|---|---|---|---|
| HCE | 100 | 113 | 285 |
| Thermo | 936 | 28039 | 18 |
| IOBA | 142 | 195 | 324 |
| MITOMICS | 430 | 113654 | 21 |
| Standards | 54 | 80379 | 4310 |
| PNNL | 1631 | 46340 | 107 |

***Table S8:*** *Number of lipid species per lipid class for each data source.*

| Lipid Class | HCE | Thermo | IOBA | MITOMICS | Standards | PNNL |
|---|---|---|---|---|---|---|
| CAR | 0 | 0 | 0 | 607 | 0 | 0 |
| CE | 0 | 0 | 0 | 275 | 0 | 64 |
| CL | 0 | 0 | 0 | 1739 | 0 | 483 |
| Cer | 16 | 358 | 2 | 12962 | 0 | 2691 |
| DG | 0 | 259 | 0 | 8 | 0 | 1674 |
| DGDG | 0 | 0 | 0 | 0 | 0 | 320 |
| EPC | 0 | 0 | 0 | 0 | 0 | 2 |
| FA | 12 | 0 | 0 | 0 | 0 | 0 |
| GalCer | 2 | 0 | 0 | 0 | 0 | 838 |
| GlcCer | 0 | 0 | 0 | 0 | 0 | 6 |
| HexCer | 3 | 123 | 4 | 0 | 0 | 0 |
| IPC | 0 | 0 | 0 | 0 | 0 | 120 |
| LPC | 0 | 0 | 0 | 9894 | 0 | 2760 |
| LPE | 1 | 861 | 0 | 22675 | 0 | 2016 |
| LPG | 0 | 26 | 0 | 2238 | 0 | 70 |
| LPI | 0 | 146 | 0 | 121 | 0 | 394 |
| LPS | 0 | 64 | 0 | 0 | 0 | 144 |
| LacCer | 1 | 0 | 0 | 0 | 0 | 66 |
| MG | 0 | 0 | 0 | 0 | 0 | 12 |
| MGDG | 0 | 0 | 0 | 0 | 0 | 42 |
| PA | 1 | 0 | 0 | 250 | 6444 | 150 |
| PC | 16 | 7565 | 72 | 22003 | 31125 | 10153 |
| PE | 27 | 2544 | 49 | 12286 | 22696 | 6730 |
| PG | 5 | 628 | 4 | 27 | 7329 | 2880 |
| PI | 8 | 2242 | 10 | 4235 | 0 | 2736 |
| PS | 6 | 2524 | 9 | 811 | 12785 | 1716 |
| SHexCer | 0 | 2 | 0 | 0 | 0 | 0 |
| SM | 15 | 28 | 7 | 0 | 0 | 1881 |
| SPB | 0 | 0 | 0 | 0 | 0 | 100 |
| SQDG | 0 | 0 | 0 | 0 | 0 | 228 |
| TG | 0 | 10669 | 38 | 14185 | 0 | 8064 |

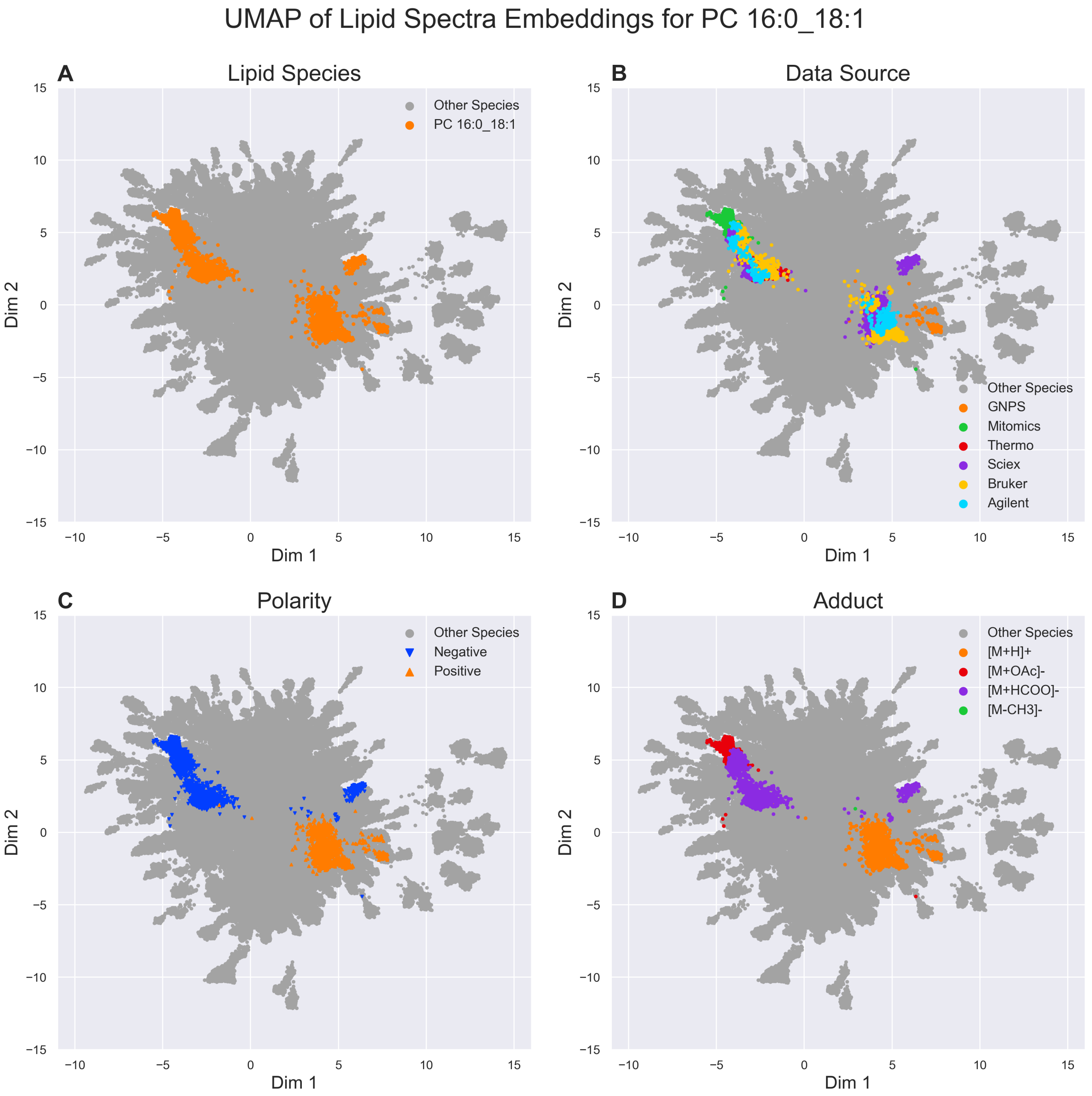


***Figure S4:*** *UMAP of spectrum embeddings for A) PC 16:0_18:1 colored by B) source, C) polarity, and D) adduct.*

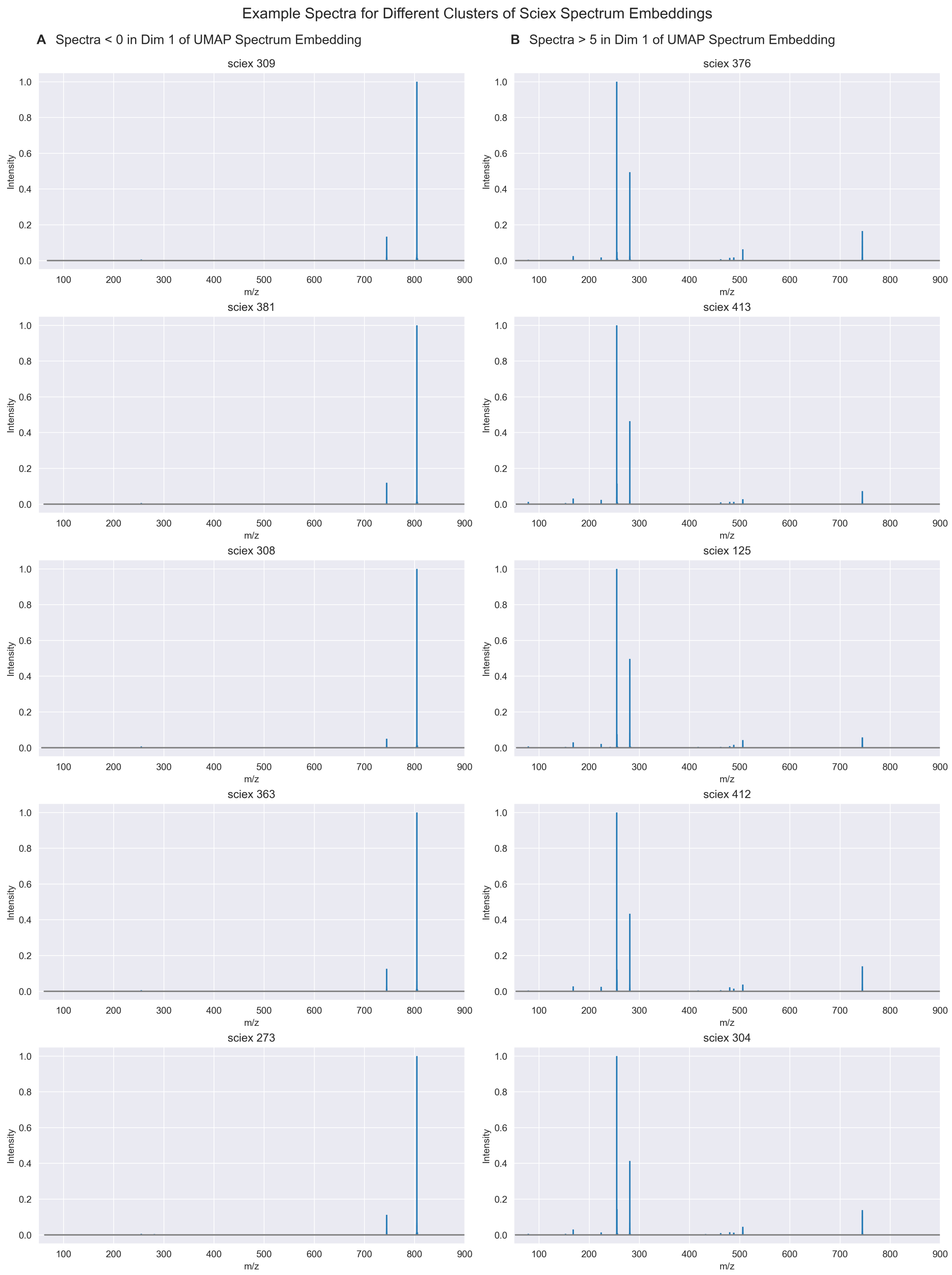


***Figure S5:*** *Example spectra for the two different sciex clusters in the UMAP embedding with A) Dim1 < 0 and B) Dim1 > 5.*

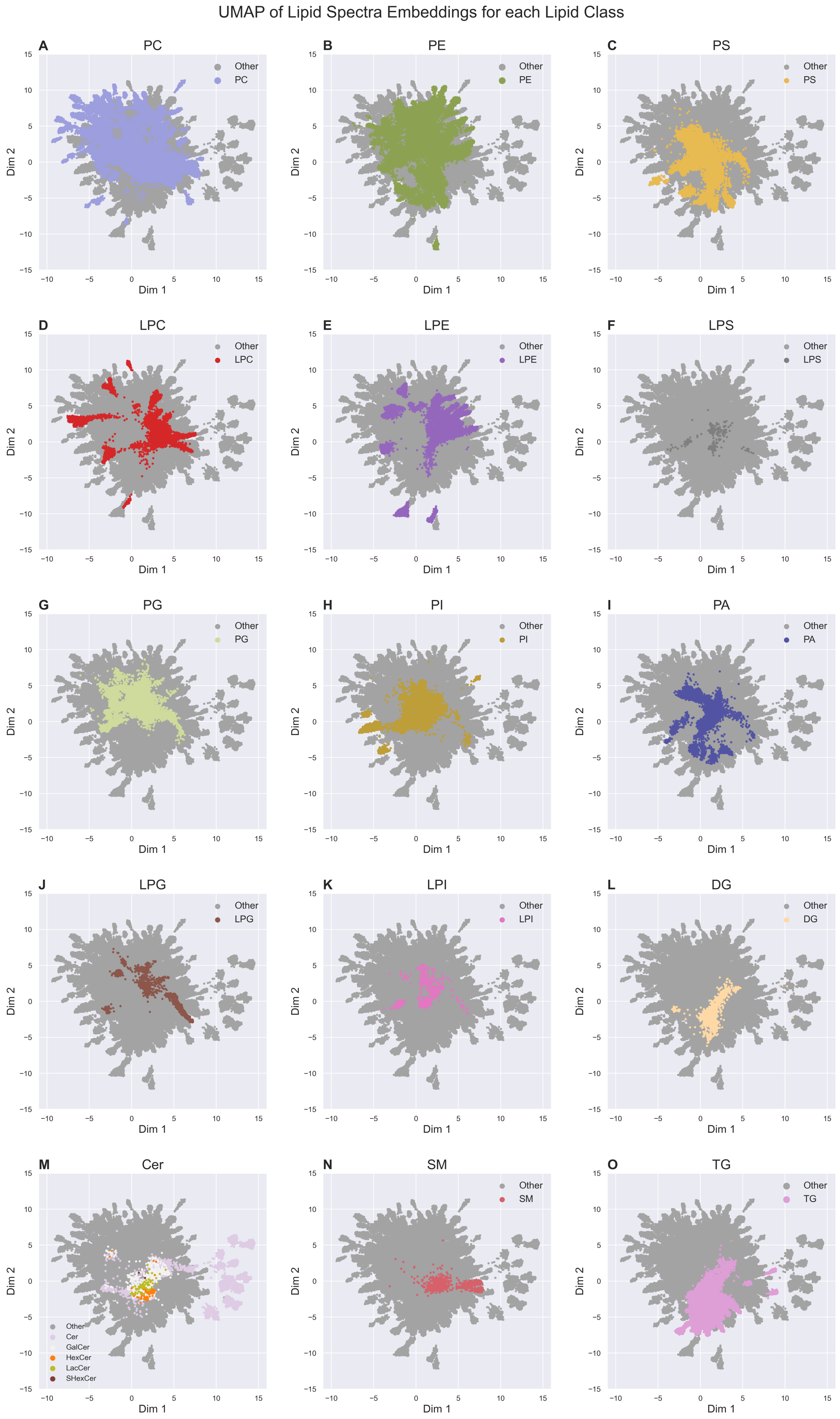


***Figure S6:*** *Umap of embeddings colored by lipid classes.*

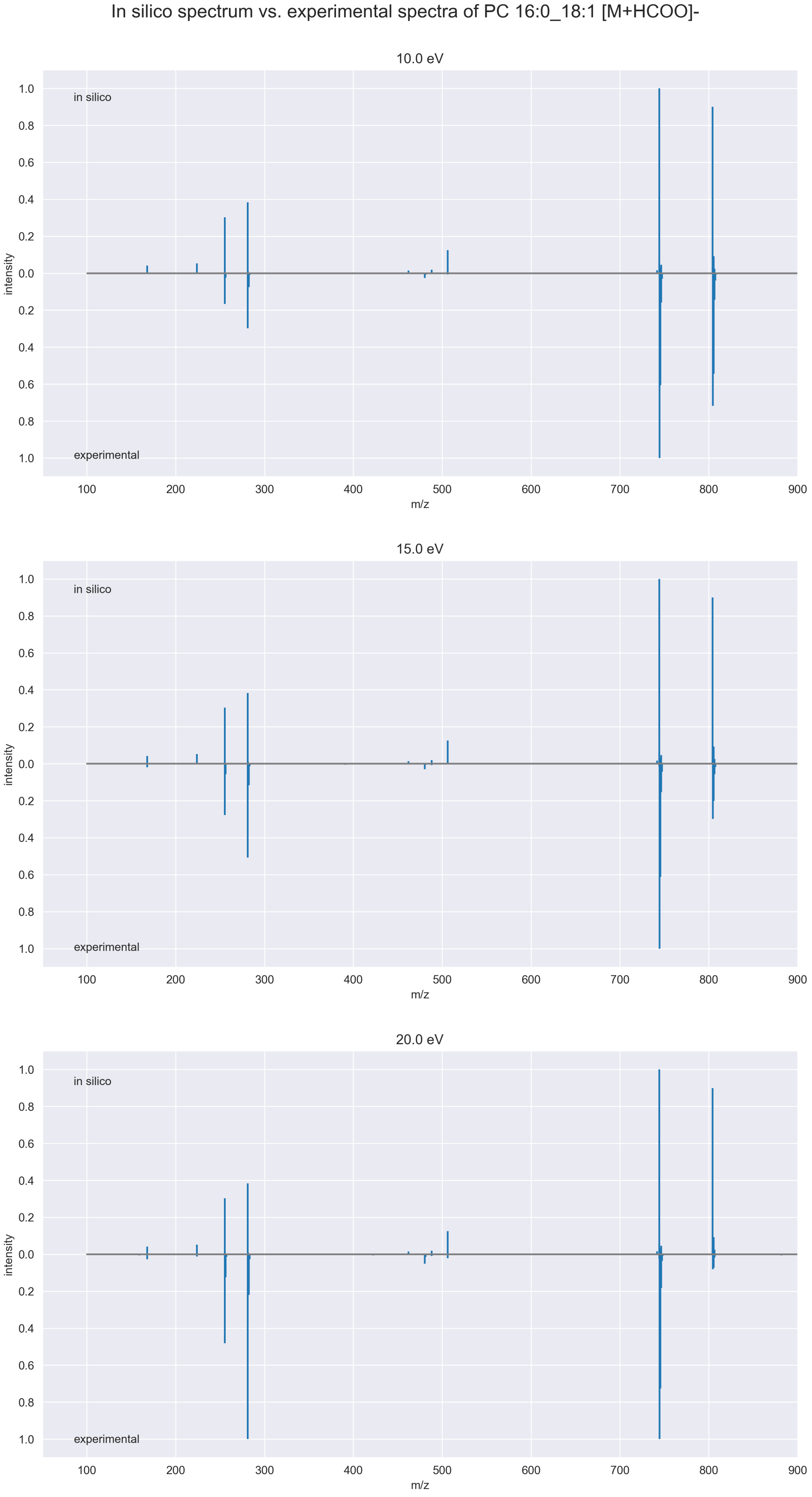


***Figure S7:*** *Comparison between predicted in silico spectrum for PC 16:0_18:1 and experimental spectra at different collision energies of A) 10 eV, B) 15 eV and C) 20 eV.*

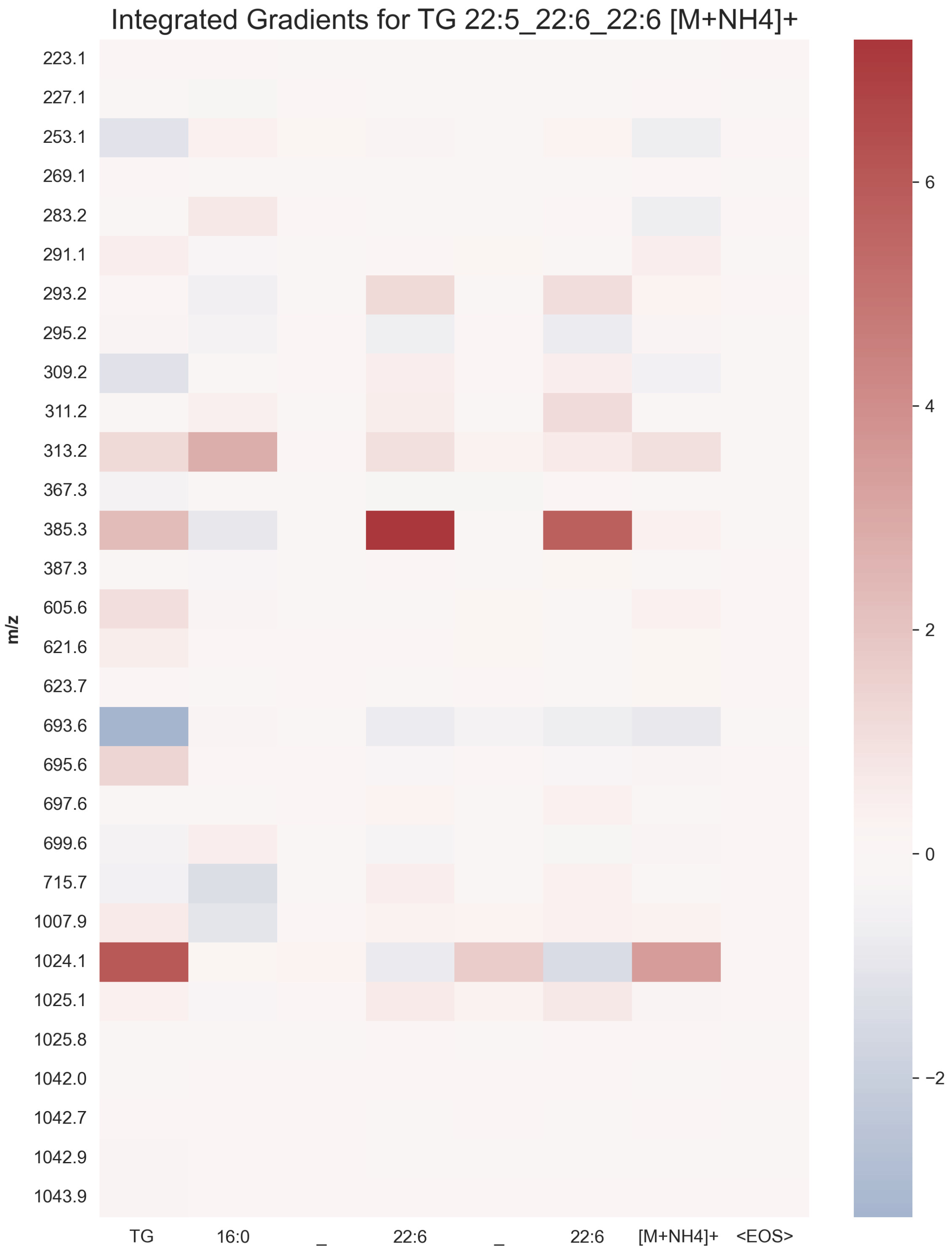


***Figure S8:*** *Integrated gradients for misprediction of true label TG 22:5_22:6_22:6 as TG 16:0_22:6_22:6.*

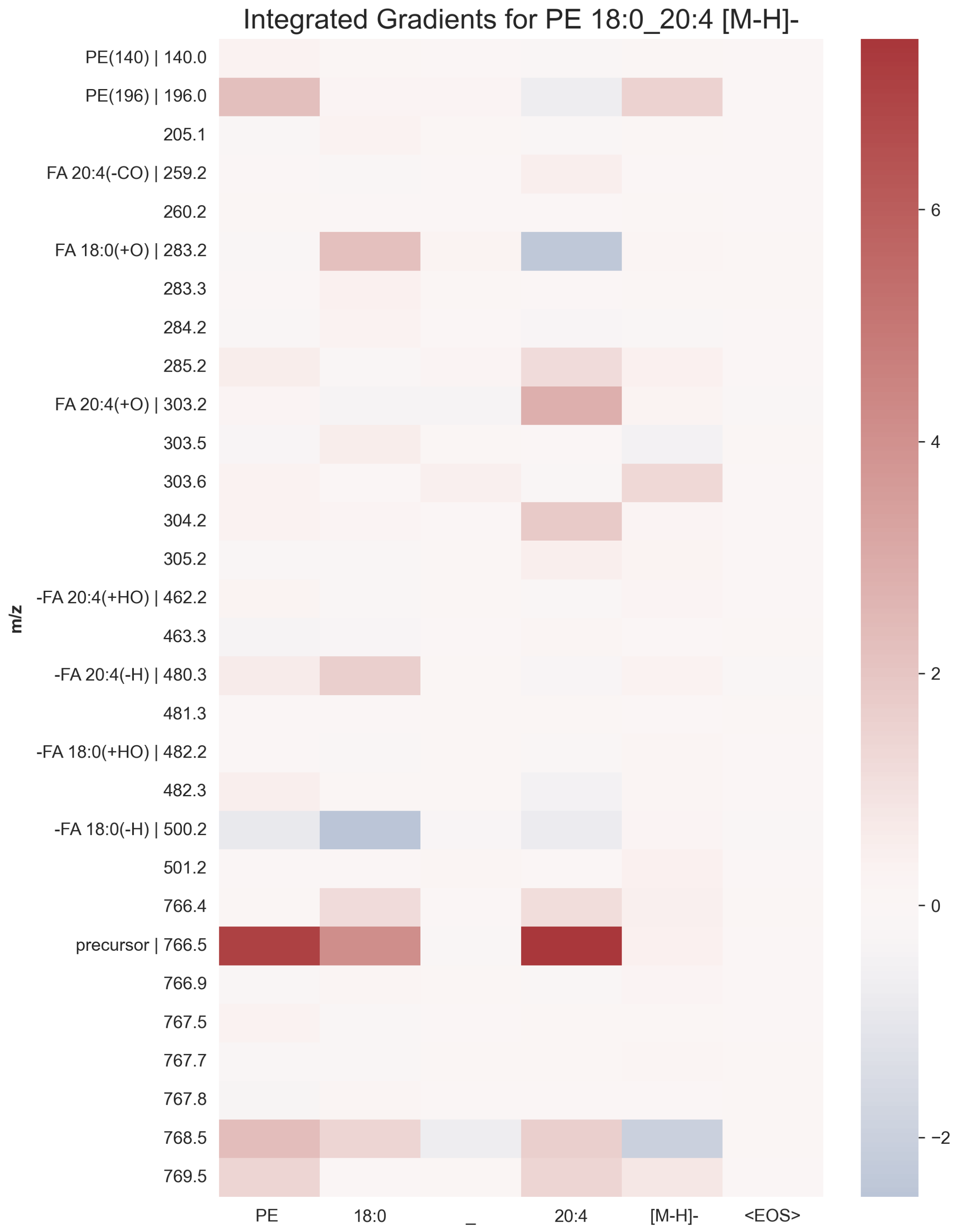


***Figure S9:*** *Example of integrated gradients for PE 18:0_20:4 [M-H]-.*

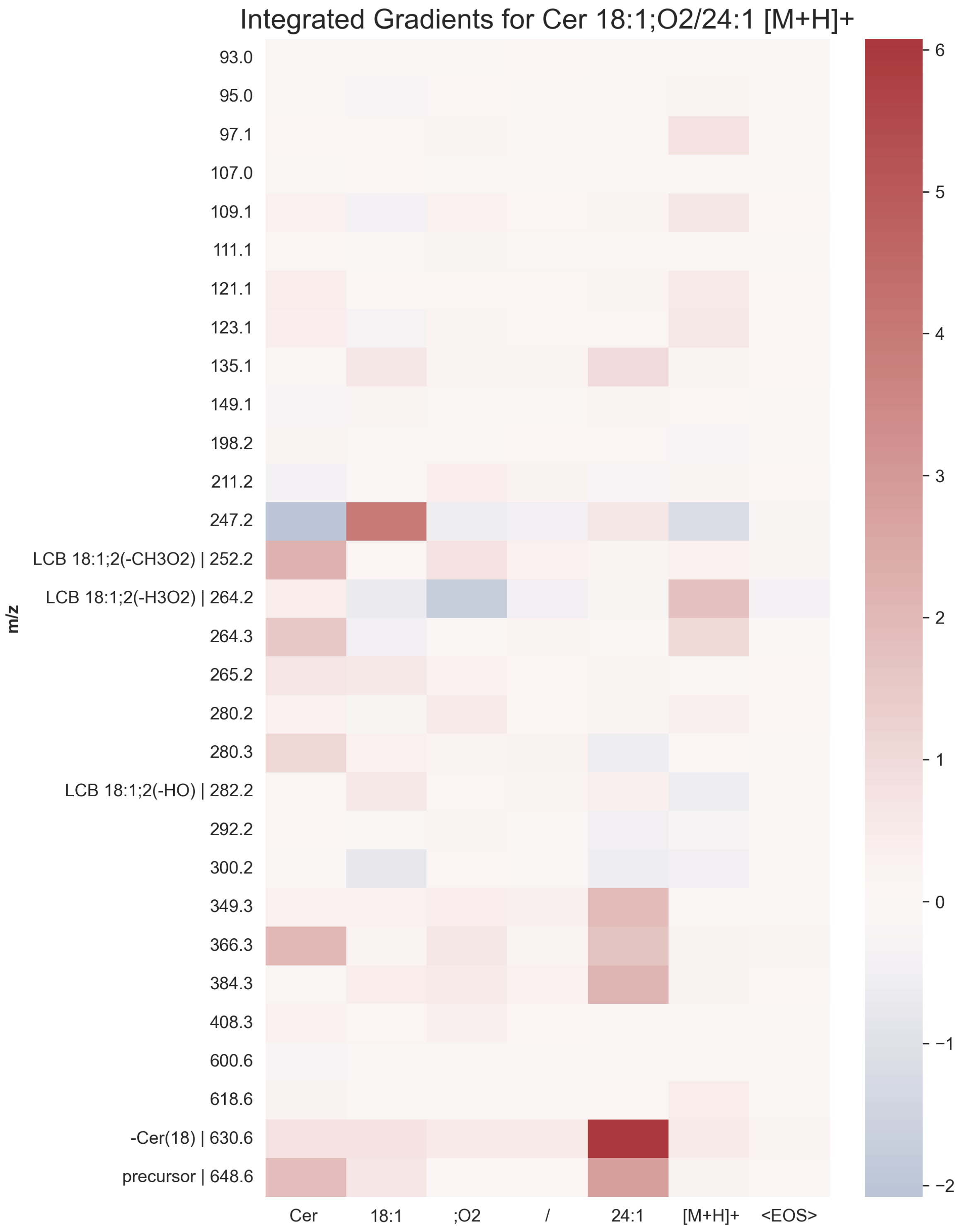


*Figure S10:* *Example of integrated gradients for Cer 18:1;O2/24:1 [M+H]+*

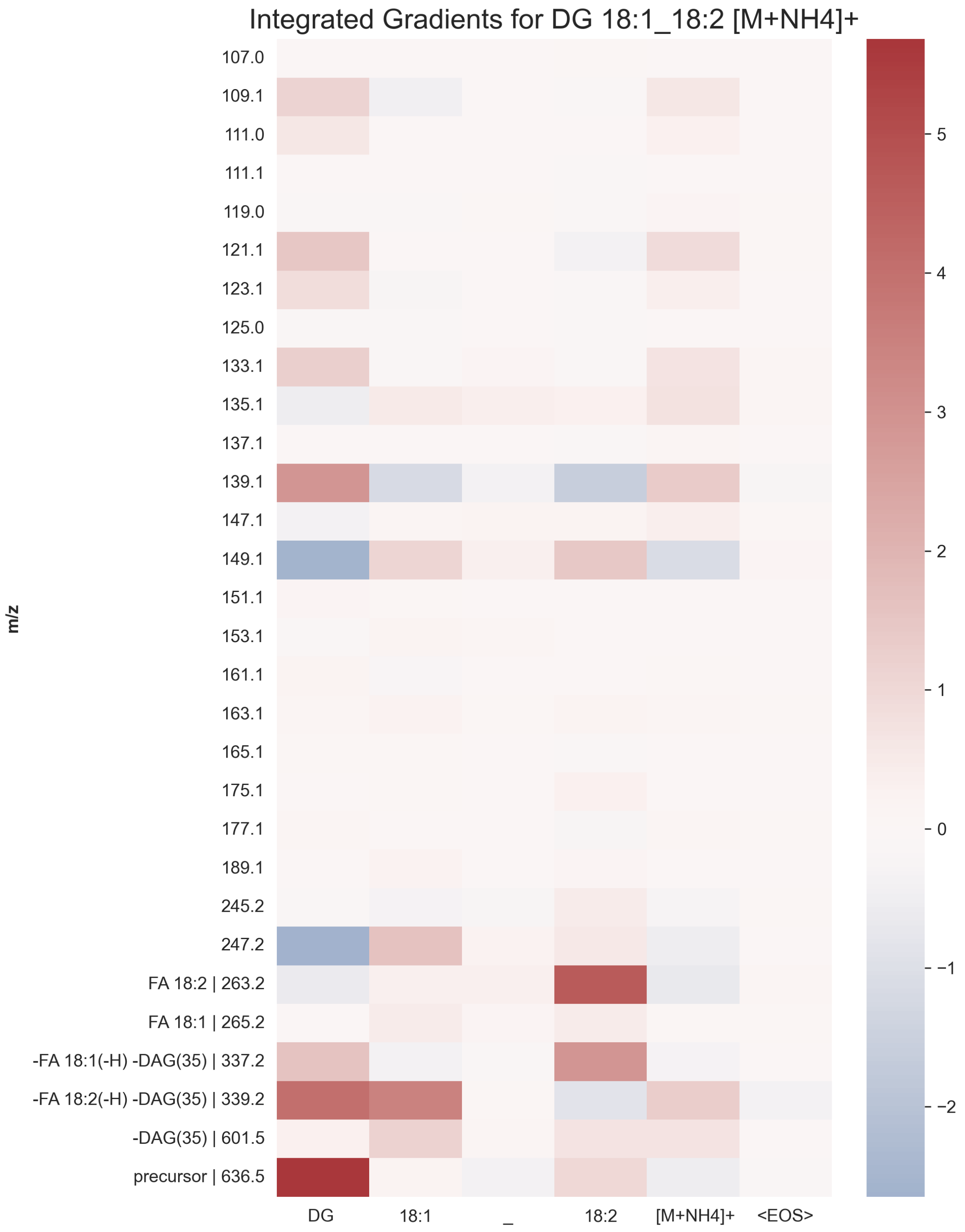


***Figure S11:*** *Example of integrated gradients for DG 18:1_18:2 [M+NH4]+*
